# New insights into the postcranial morphology of *Lithornis vulturinus* from the Eocene London Clay

**DOI:** 10.64898/2026.03.17.711321

**Authors:** Klara Widrig, Daniel J. Field

## Abstract

The deepest phylogenetic divergence within crown birds (Neornithes) is that between the reciprocally monophyletic Palaeognathae and Neognathae. Extant palaeognath diversity comprises the iconic flightless “ratites” (ostriches, rhea, kiwi, cassowaries, and emu), as well as 46 species of volant tinamous in Central and South America (Billerman et al., 2020). Although the earliest stages of palaeognath evolution remain shrouded in mystery due to a sparse fossil record, a group of apparently volant extinct palaeognaths from the Paleogene of Europe and North America, the lithornithids, can help to clarify palaeognath origins. Here, we use high resolution microCT scanning to characterize the morphology of two lithornithid specimens from the early Eocene (Ypresian) London Clay Formation: the neotype of *Lithornis vulturinus* (NHMUK A5204), from the Isle of Sheppey, Kent, England, and a newly discovered clay nodule containing lithornithid postcranial remains from the nearby locality of Seasalter. This three-dimensional dataset reveals bones from the *L. vulturinus* neotype that are partially or completely covered by matrix, allowing us to redescribe this critical specimen in new detail and present a revised differential diagnosis of *L. vulturinus*. We refer the new specimen from Seasalter to *L. vulturinus* on the basis of apomorphies such as a proximally directed lateral process of the coracoid, caudally divergent lateral margins of the sternum, an arcuate deltopectoral crest, as well as its provenance from a nearby penecontemporaneous locality. The Seasalter specimen contains abundant postcranial material that provides new insight into bones damaged or missing in the neotype, including two undamaged scapulae bearing the hooked acromion that is a diagnostic feature of lithornithids, two complete coracoids, and a nearly complete three-dimensionally preserved sternum. Its estimated body mass is one third larger than that of the neotype, indicating intraspecific variation within *L. vulturinus* that may reflect sexual dimorphism. Molecular divergence dates and Cretaceous neognath fossils indicate the presence of total-clade palaeognaths before the K–Pg mass extinction event; detailed anatomical descriptions of Paleogene palaeognaths will assist in the identification of the first total-clade palaeognaths from the Cretaceous, and provide insight into how and when flight was independently lost among Cenozoic crown palaeognaths.

## Introduction

Lithornithids, a group of palaeognathous birds known from the middle Paleocene to the early Eocene in North America (Houde, 1988, Stidham et al., 2014) and the middle Paleocene to the middle Eocene in Europe (Mayr, 2008, Mayr, 2009b, Mayr and Smith, 2019), include one of the first extinct bird taxa to be described scientifically (Houde, 1988). Found on the Isle of Sheppey in southeastern England by the anatomist John Hunter, the holotype specimen of the bird that would later become known as *Lithornis vulturinus* was purchased from his collections following his death in 1793 by the Royal College of Surgeons in 1798 (Houde, 1988, Leonard et al., 2005). This holotype consisted of a clay nodule with an embedded partial sternum, thoracic vertebra, ribs, the distal end of a left femur, and a proximal end of a left tibiotarsus (Houde, 1988). It was not published on until decades later by Owen (1840) in one of the first publications to treat an extinct bird, where it was interpreted as a cathartid vulture (Accipitriformes: Cathartidae) on the basis of the similarity in shape of intermuscular lines on the coracoid and sternum, as well as “the same general form and proportions of the bones”. It was figured by Owen (1841) in two lithographs and as a woodcut in 1846. Sadly, this specimen was destroyed in the bombing of London during the Second World War and is now known only from the woodcut and lithographs (Harrison and Walker, 1977, Houde, 1988).

Given the unfortunate loss of this scientifically and historically significant fossil, Houde (1988) designated another specimen, NHMUK A5204, as the neotype for *L. vulturinus*. NHMUK A5204 is a phosphate nodule and associated fragments collected by J. Quayle from Warden Point on the Isle of Sheppey (Wharton, 2002) and donated to the UK Natural History Museum in 1979. Houde described the specimen as containing a right humerus, radius, and ulna each lacking their distal ends, a right scapula and the right half of the sternum, the distal ends of a left radius and ulna, the proximal portions of a left femur and right tibiotarsus, vertebrae C10, C12, C13, C15, and T1, ribs, and various bone fragments. A partial neurocranium separate from the main phosphatic nodule but also designated NHMUK A5204 was not described by Houde (1988). The earliest mention of this neurocranium in the literature is in an unpublished doctoral thesis (Wharton, 2002). We treat this neurocranium in a separate publication (Widrig et al., 2024). Despite sharing few elements with the destroyed holotype, Houde (1988) was confident in his assignment of the neotype, as referred specimens that share elements with NHMUK A5204 and the lost holotype show they are consistent in morphology.

Additional fossil discoveries including palatal material had already enabled the recognition of lithornithids as palaeognaths by the time of Houde’s 1988 monograph, with representatives of lithornithids known from both sides of the Atlantic (Houde and Olson, 1981, Houde, 1986, Houde, 1988). *Lithornis vulturinus*, *Lithornis nasi*, ?*Lithornis hookeri*, *Fissuravis weigelti*, ?*Pseudocrypturus danielsi*, and ?*Pseudocrypturus gracilipes* represent the named species from Europe (Houde, 1988, Mayr, 2007, Mayr and Kitchener, 2025). Of these, *L. nasi* has been considered a junior synonym of *L. vulturinus* (Bourdon and Lindow, 2015), and very little material exists for ?*Lithornis hookeri*, as one may surmise by its uncertain placement in this genus (Houde, 1988). *Fissuravis*, known only from a partial coracoid, lacks clear diagnostic features and appears to be missing a foramen on the posteroventral surface of the acrocoracoid process that is diagnostic of Lithornithidae (Nesbitt and Clarke, 2016). In North America, lithornithids are thus far represented by six species in four genera. The lithornithid affinities of these birds, *Lithornis celetius*, *Lithornis promiscuus*, *Lithornis plebius*, *Paracathartes howardae*, *Pseudocrypturus cercanaxius*, and *Calciavis grandei* (Houde, 1988, Nesbitt and Clarke, 2016), have not been questioned. Fossils tentatively assigned to *Pseudocrypturus cercanaxius* and *Calciavis grandei* (*Lithornis* cf. *grandei* sensu Mayr and Kitchener, 2025) from the London Clay Formation of England, and *Lithornis nasi* from the Willwood Formation of Wyoming, USA (Houde, 1988) suggest that some species may have exhibited geographic distributions spanning the Atlantic Ocean.

The question of whether these five genera comprise a monophyletic group, a paraphyletic grade of stem palaeognaths, or even a polyphyletic group in which lithornithids represent stem members of different clades within Palaeognathae, has not been unambiguously resolved (Nesbitt and Clarke, 2016, Widrig and Field, 2022). Houde (1988) argued against a monophyletic Lithornithidae and placed *Paracathartes* closer to extant palaeognaths than to other lithornithids on the basis of bone histology he considered more similar to that of ‘ratites’ than to other lithornithids, though the histological difference between *Paracathartes* and other lithornithid genera may in part be due to the larger body size of *Paracathartes* relative to other lithornithids (Mayr, 2009a). Nesbitt and Clarke (2016) found strong support for a monophyletic Lithornithidae, but were unable to achieve any resolution within the clade. A monophyletic Lithornithidae is supported by most recent analyses (Worthy et al., 2016, Worthy et al., 2017, Yonezawa et al., 2017) with the exception of Yonezawa et al. (2017)’s alternative maximum likelihood analysis using ten non-homoplastic characters from Houde (1988), which placed *Pseudocrypturus* as a stem palaeognath outside a *Lithorni*s + *Paracathartes* clade and rendered Lithornithidae a paraphyletic grade.

Lithornithids are generally accepted to be total-clade palaeognaths, but their position within Pan-Palaeognathae is dependent on the prior constraints applied in phylogenetic analyses. When trees are constrained to match recent molecular phylogenetic topologies (in which ratites are paraphyletic with respect to tinamous), lithornithids are frequently recovered as stem palaeognaths, but otherwise are recovered as sister to tinamids likely due to the highly derived morphology of the remainder of extant palaeognaths, the flightless ‘ratites’ (Nesbitt and Clarke, 2016, Yonezawa et al., 2017, Almeida et al., 2022). Regardless of their position within Pan-Palaeognathae, lithornithids, like all other palaeognaths, exhibit a palate characterized by fused pterygoids and palatines in articulation with enlarged basipterygoid processes, a single articular facet for the otic capitulum of the quadrate, a dentary that is deeply forked caudally, and open ilioischiadic foramina (Pycraft, 1900, Bock, 1962, Parkes and Clark, 1966, Cracraft, 1974, Houde, 1986, Mayr and Zelenkov, 2021, Widrig and Field, 2022, Crane et al., 2025, Plateau et al., 2026). With bills that appear capable of distal rhynchokinesis and evidence for a functional vibrotactile bill tip probing organ, some authors have suggested lithornithids were ground-foraging probe feeders functionally akin to ibises or kiwi (Houde, 1988, du Toit et al., 2020). However, unlike many ground-dwelling birds whose flight abilities are limited mainly to short bursts to escape predators (e.g. Tinamidae, Phasianidae) or which have lost flight entirely, lithornithids appear to have been capable of long-distance flight (Houde, 1988, Torres et al., 2020, Widrig and Field, 2022, Widrig et al., 2025). This potential capacity of lithornithids to disperse over wide areas has led to renewed interest in light of molecular phylogenetic hypotheses that show the volant tinamous nested within the flightless ‘ratites’ (Hackett et al., 2008, Harshman et al., 2008, Phillips et al., 2009, Haddrath and Baker, 2012, Smith et al., 2012, Baker et al., 2014, Mitchell et al., 2014, Claramunt and Cracraft, 2015, Prum et al., 2015, Grealy et al., 2017, Reddy et al., 2017, Yonezawa et al., 2017, Cloutier et al., 2019, Kimball et al., 2019, Sackton et al., 2019, Feng et al., 2020, Kuhl et al., 2020, Urantówka et al., 2020, Almeida et al., 2022, Wang et al., 2021, Takezaki, 2023, Stiller et al., 2024), upending our understanding of palaeognath interrelationships and biogeography from one consistent with vicariant speciation due to continental drift (Cracraft, 1973, Cracraft, 1974, Roff, 1994, Cracraft, 2001) to a more complex scenario in which a presumably volant ancestral group underwent multiple overseas dispersal events followed by independent losses of flight.

Looking further stemward, there exists substantial molecular evidence for a Cretaceous divergence between the reciprocally monophyletic palaeognaths and neognaths (Phillips et al., 2009, Haddrath and Baker, 2012, Jarvis et al., 2014, Mitchell et al., 2014, Claramunt and Cracraft, 2015, Prum et al., 2015, Grealy et al., 2017, Yonezawa et al., 2017, Sackton et al., 2019, Kuhl et al., 2020, Almeida et al., 2022, Wang et al., 2021, Stiller et al., 2024). Given the very well-supported sister group relationship between palaeognaths and neognaths, this preponderance of molecular evidence as well as fossils likely belonging to neognaths from the latest Cretaceous (Clarke et al., 2005, Field et al., 2020, Torres et al., 2025, Irazoqui et al., 2026) implies that palaeognaths, possibly including lithornithids, should have existed alongside them. However, no definitive palaeognath fossils from the Mesozoic have been identified. A partial scapula from the Hornerstown Formation in New Jersey, USA could extend the temporal range of lithornithids back as early as the latest Maastrictian/earliest Danian (Parris and Hope, 2002). The scapula was attributed to a lithornithid due to its hooked acromion process, but this is not sufficient for a reliable referral as several Mesozoic non-neornithine ornithurines possess a hooked acromion approaching the condition seen in lithornithids (Clarke, 2004, Nesbitt and Clarke, 2016, Benito et al., 2022a). Further fossil discoveries are obviously needed, but more detailed descriptions of lithornithid anatomy could assist in recognizing older palaeognath fossils that may have been overlooked. Given that evidence has emerged suggesting that the neornithine most recent common ancestor had a neognath-like palate (Benito et al., 2022b), postcranial anatomy could be of particular importance for recognizing the earliest palaeognaths, as the characteristic palaeognathous palate may have taken longer to evolve.

Lithornithid morphology has been treated in detail in recent years. For instance, an exquisitely preserved specimen of *L. vulturinus* from the Fur Formation of Denmark has been thoroughly described (Leonard et al., 2005, Bourdon and Lindow, 2015), and Nesbitt and Clarke (2016) provide an in-depth description of *Calciavis grandei* from the Green River Formation. Most recently, Mayr and Kitchener (2025) describe the lithornithids donated by Michael Daniels to the National Museums Scotland from Walton-on-the-Naze, Essex, UK. Meanwhile, the *L. vulturinus* neotype has eluded reinvestigation since Houde’s 1988 monograph, in which the skeletal elements present in NHMUK A5204 were listed. Houde (1988) diagnosed *L. vulturinus* as simply being smaller than *L. promiscuus* but larger than the remainder of its congeners *L. nasi,* ?*L. hookeri, L. celetius,* and *L. plebius*, with a more arcuate deltopectoral crest and more caudally divergent margins of the sternum than in other members of the genus.

In this study, we provide a detailed redescription and differential diagnosis of the neotype of *Lithornis vulturinus* (Figure 1). Given that *Lithornis vulturinus* is the type species for the genus *Lithornis*, which is in turn the type genus for the clade Lithornithidae (Houde, 1988), we aim to provide a timely update in light of the increased interest surrounding this clade. Additionally, we present a new specimen of the same species found near the town of Seasalter (Figure 1), only a few kilometres away from the type locality on the Isle of Sheppey. We use high resolution microCT scanning to reveal bones that were partially or completely obscured by matrix in the neotype, and to digitally extract the Seasalter partial skeleton from the phosphate nodule it was found in. This method has previously been successfully applied to London Clay fossils still within matrix (Beckett et al., 2016). We hope that these anatomical descriptions will aid in both clarifying the species limits and systematics of Lithornithidae as well as the identification of palaeognaths from the Late Cretaceous and early Paleocene.

**Figure 1.**
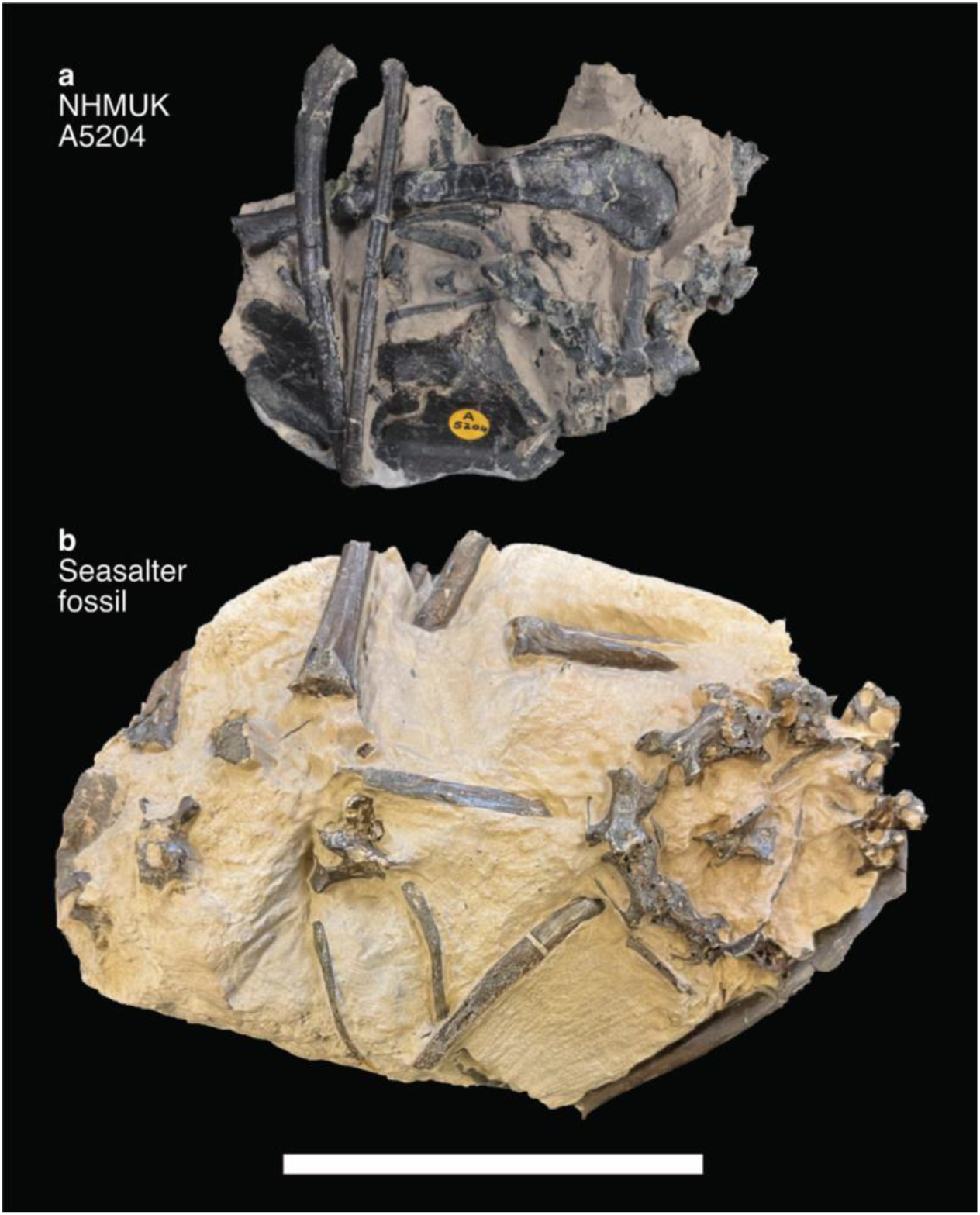
Photographs of specimens described in this publication. a) The prepared nodule containing the neotype specimen NHMUK A5204, and b) the partially prepared nodule containing the Seasalter specimen. Scale bar = 10 cm.

## Methods

The neotype was scanned on a Nikon Metrology XT H 225 ST High Resolution CT Scanner located at the Natural History Museum, London. Scanning parameters were: 175 kV, 257 uA, 3841 projections, 85 frames, 8 frames per projection. The new fossil from Seasalter was scanned at the Cambridge Biotomography Centre on a Nikon Metrology XT H 225 ST High Resolution CT Scanner. Scanning parameters were: 180 kV, 250 uA, 3142 projections. All scanned material was digitally segmented and rendered in VGSTUDIO MAX version 3.4 (Volume Graphics, Heidelberg, Germany). 3D surface meshes of each element were extracted and exported using this program.

We refer primarily to Houde (1988) for comparative morphological information on lithornithids, with additional information from Leonard et al. (2005), Bourdon and Lindow (2015), Nesbitt and Clarke (2016), and Mayr and Kitchener (2025).

Comparisons with the outgroup taxon *Ichthyornis dispar* were based on Benito et al. (2022a). We based our comparisons with extant taxa on specimens from the University of Cambridge Museum of Zoology (UMZC), particularly on the Elegant-Crested Tinamou *Eudromia elegans* specimen UMZC 404.E, as well as the character states of tinamids described by Bertelli (2017). Anatomical terms follow Baumel and Witmer (1993) unless otherwise stated, with English equivalents used when applicable.

We tested the phylogenetic positions of the neotype and Seasalter specimen by scoring the two fossils into the morphological data matrix of Nesbitt and Clarke (2016). This matrix contains 182 characters scored for 38 terminal taxa. Characters in this matrix were derived from the morphological datasets of Cracraft (1974), Bledsoe (1988), Lee et al. (1997), Mayr and Clarke (2003), Clarke (2004), and Clarke et al. (2006), as well as new observations by the authors. Though it was included by Nesbitt and Clarke (2016), we chose to remove *Apsaravis ukhaana* from our analyses, as it has been identified as a wildcard taxon in several analyses, e.g.,Longrich et al. (2011), Field et al. (2018), and Benito et al. (2022a). Several lithornithid and extant palaeognath character states were amended based on the observations of Mayr and Kitchener (2025). We removed characters 105 and 164 from the Nesbitt and Clarke (2016) matrix as we found them to be ambiguous.

We performed separate phylogenetic analyses using parsimony and Bayesian analytical frameworks. We conducted maximum parsimony analyses using TNT version 1.5 (Goloboff and Catalano, 2016), made available with the sponsorship of the Willi Henning Society. For both our unconstrained analyses and analyses constrained to a molecular phylogenetic backbone, we performed an unconstrained heuristic search with equally weighted characters. 1,000 replicates of random stepwise addition were produced using the tree bisection reconnection (TBR) algorithm, and 10 trees were saved for each replicate. *Ichthyornis dispar* was designated as the outgroup taxon. All most parsimonious trees (MPTs) were used to calculate a strict consensus, and bootstrap values were calculated using a traditional search with 1,000 replicates, with the outputs saved as absolute frequencies.

We conducted Bayesian analyses following the protocol of Field et al. (2020) and Benito et al. (2022a) in MrBayes (Ronquist et al., 2012) using the CIPRES Science Gateway (Miller et al., 2010). Data were analysed under the Mkv model (Lewis, 2001). We assumed gamma-distributed rate variation to allow for variation in evolutionary rate across different characters. We conducted the analyses using four chains and two independent runs, with a tree sampled every 4,000 generations and a burn-in of 25%. We ran the analyses for 30,000,000 generations. We used the standard diagnostics provided in MrBayes to assess analytical convergence (average standard deviation of split frequencies <0.02, potential scale reduction factors = 1, effective sample sizes > 200). Using the sump and sumt commands, we obtained results from independent runs of the same analyses. We exported the recovered trees into TNT to optimize morphological synapomorphies of recovered tree topologies.

We repeated our maximum parsimony and Bayesian analyses with phylogenetic constraints based on molecular phylogenetic topologies. Palaeognath relationships were constrained to match the molecular topologies of Phillips et al. (2009), Mitchell et al. (2014), Grealy et al. (2017), Yonezawa et al. (2017), Urantówka et al. (2020), Almeida et al. (2022), and the concatenated dataset of Cloutier et al. (2019). Neognath relationships were constrained to match those of Prum et al. (2015). For our final set of maximum parsimony and Bayesian analyses, lithornithids were additionally constrained as the sister taxon of all other palaeognaths.

Lithornithid body masses were estimated using the equations of Field et al. (2013). All body size measurements of the neotype and new block were obtained in VGSTUDIO MAX version 3.4 using the program’s “measure” function. Relevant measurements of Walton on the Naze specimens housed at the National Museums Scotland (NMS) were taken from Mayr and Kitchener (2025), those for MGUH 26770 were obtained from Bourdon and Lindow (2015), and those for SGPIMH MEV1 were obtained from Mayr (2009b). All other measurements were obtained from specimens in the collections of the US National Museum of Natural History (USNM) using Mitutoyo Absolute Digimatic callipers accurate to the nearest 0.01 mm, and from the UK National Museum of Natural History (NHMUK) using Fisher brand callipers accurate to the nearest 0.2 mm. Where two measurements were available for the same individual due to both left and right elements being preserved, the resulting body mass estimates for that element were averaged.

### Geological setting (Figure 2)

Both the holotype, neotype, and new specimen of *Lithornis vulturinus* discussed herein derive from the London Clay Formation, a marine deposit that consists primarily of heavily bioturbated argillaceous or slightly calcareous clay that crops out over a wide area in southeastern England (Friedman et al., 2016). Fossil fish are extremely abundant in this formation, though terrestrial fossils including plants, insects, and vertebrates (including the birds discussed here) appear to have been transported offshore and fossilized there (Friedman et al., 2016). Typically, vertebrates in the London Clay are found preserved within phosphatic or calcitic nodules (Allison, 1988).

**Figure 2.**
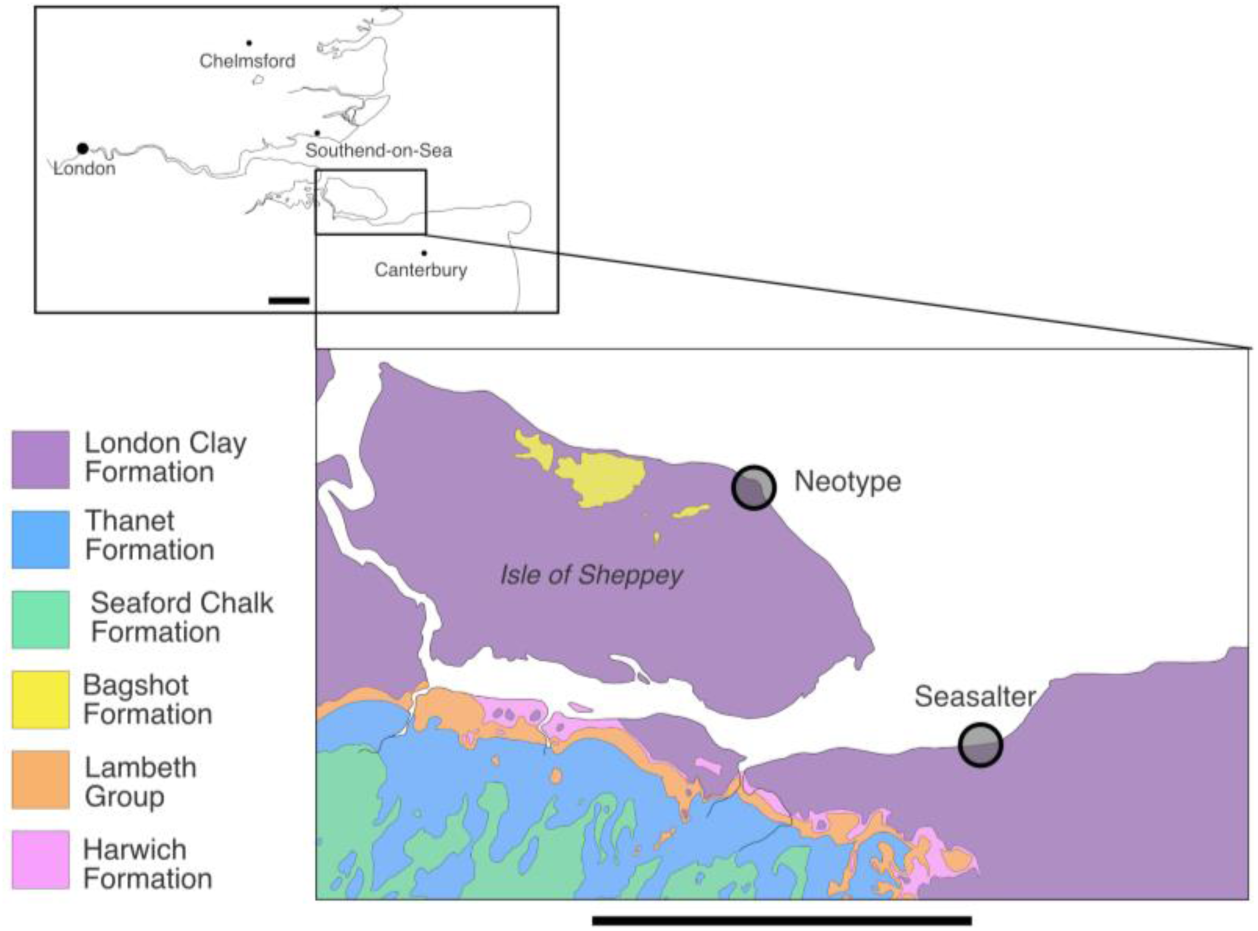
Geologic map displaying the approximate localities of the neotype (NHMUK A5204) and the new Seasalter specimen in southeast England, indicated with open circles. The London Clay Formation is shown in purple, the Thanet Formation in blue, the Seaford Chalk Formation in green, the Bagshot Formation in yellow, the Lambeth Group in orange, and the Harwich Formation in pink. Scale bars = 10 km. Geologic map inset redrawn from the British Geological Survey Geology Viewer bedrock map, containing British Geological Survey materials © UKRI [2024] (www.bgs.ac.uk/map-viewers/bgs-geology-viewer/).

The London Clay Formation is subdivided into divisions A through F, with divisions C through F found on the Isle of Sheppey (King, 1981, King, 1984, King, 2016, Collinson et al., 2016). According to the framework of Vandenberghe et al. (2012), Isle of Sheppey fossils fall between 53.5 and 51.5 Ma, placing them within the Ypresian Stage of the early Eocene. As the London Clay subdivision from which the neotype derived is unknown, no further age precision is possible.

### Institutional Abbreviations

MGUH: Natural History Museum of Denmark, Copenhagen, Denmark; NHMUK: National Museum of Natural History, London, UK; NMS: National Museums Scotland, Edinburgh, UK; SGPIMH: Geologisch-Paläontologisches Institut und Museum der Universität Hamburg, Hamburg, Germany; UMZC: University of Cambridge Museum of Zoology, Cambridge, UK; USNM: Smithsonian National Museum of Natural History, Washington D.C., USA.

### Anatomical Description: Neotype

#### Vertebral column (Figures 3, 4, 5)

A total of twelve vertebrae are preserved. Houde (1988) identified C10, C12, C13, C15, and T1. Several more vertebrae were completely encased in matrix and are described here for the first time. With the exception of C14, C15, and T1, the vertebral column is completely disarticulated. The vertebrae are heterocoelous. Vertebrae were identified via comparison with MGUH 26770, which preserves a complete articulated cervical and thoracic vertebral column (Leonard et al., 2005, Bourdon and Lindow, 2015), and Houde (1988)’s original description.

**Figure 3.**
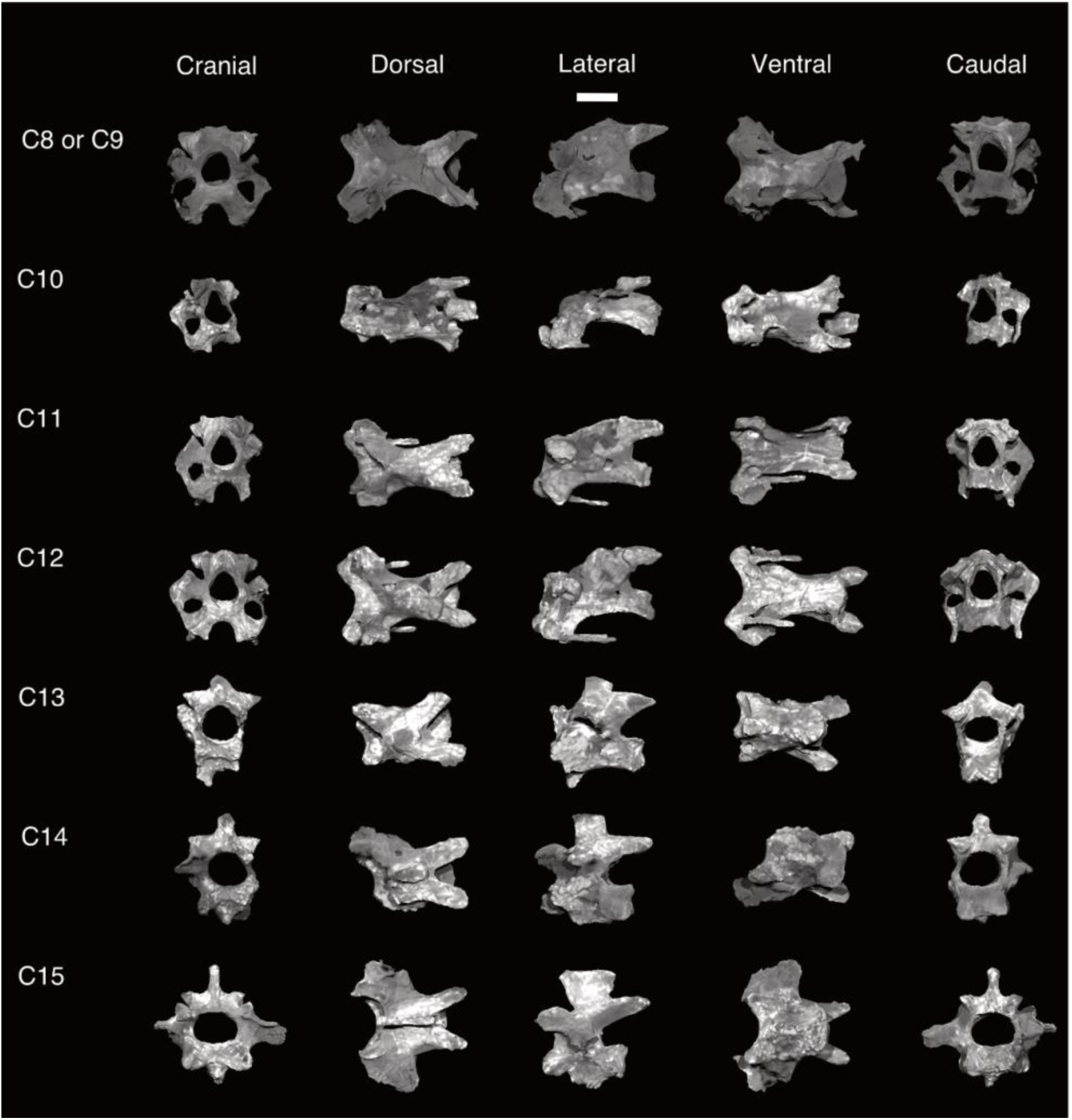
Cervical vertebrae of *Lithornis vulturinus* NHMUK A5204. Possible 8^th^ or 9^th^ cervical vertebra, possible 10^th^ through 15^th^ cervical vertebrae; in cranial, dorsal, lateral, ventral, and caudal views. Scale bar equals 5 mm.

**Figure 4.**
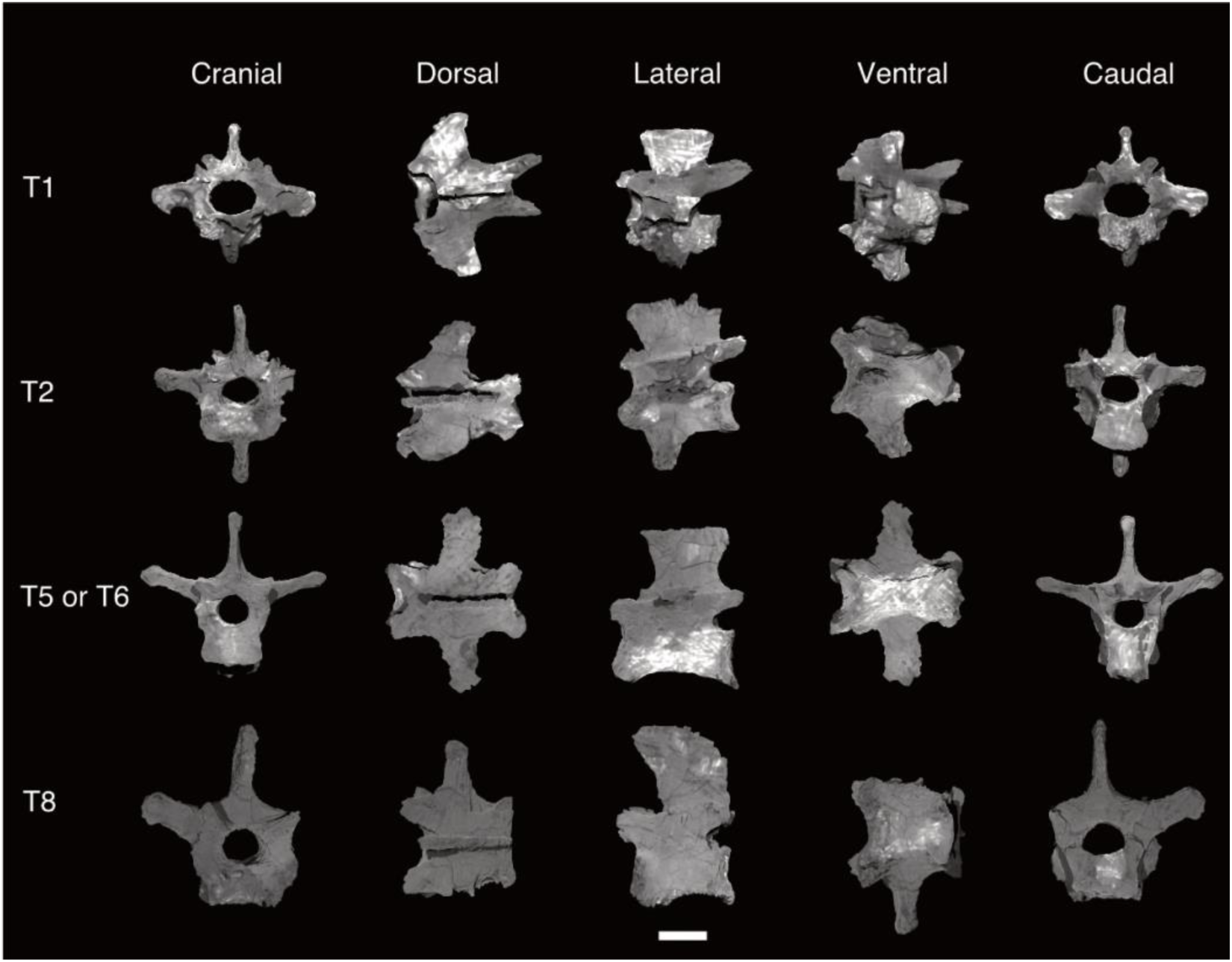
Thoracic vertebrae of *Lithornis vulturinus* NHMUK A5204. 1^st^ and 2^nd^ thoracic vertebrae, 5^th^ or 6^th^ thoracic vertebra; in cranial, dorsal, lateral, ventral, and caudal views. Scale bar equals 5 mm.

**Figure 5.**
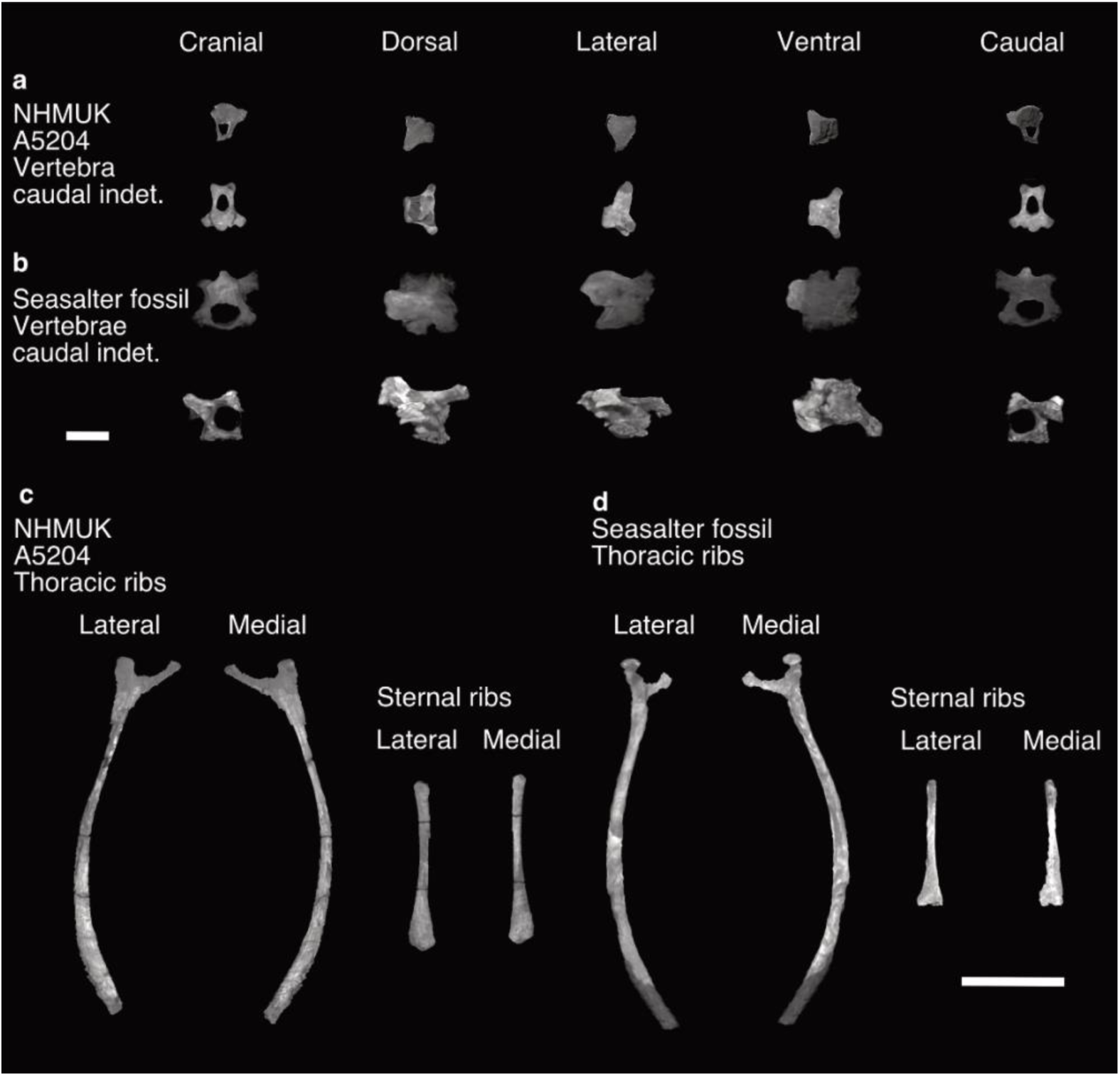
Caudal vertebrae and ribs of *Lithornis vulturinus*. (a) NHMUK A5204, caudal vertebrae indeterminate, (b) Seasalter fossil, caudal vertebrae indeterminate; in cranial, dorsal, lateral, ventral, and caudal views. Scale bar equals 5 mm. (c) NHMUK A5205, thoracic and sternal ribs, (d) Seasalter fossil, thoracic ribs and sternal ribs; in lateral and medial views. Scale bar equals 15 mm.

### Cervical vertebrae (Figure 3)

Bourdon and Lindow (2015) reported a total of 15 cervical vertebrae for *L. vulturinus* based on their observations of MGUH 26770, with the atlas, axis, and C3 to C5 comprising the cranial series, C6 to C12 the intermediate series, and C13 to C15 the caudal series as described by Baumel and Witmer (1993). Of these 15 vertebrae, an intermediate cervical vertebra most likely representing C8 or C9, and C10 through C15 are preserved in the neotype.

When preserved, the zygapophyses of the intermediate and caudal cervical vertebrae are slender, and longer than they are wide. The transverse processes are small and do not project far laterally from the vertebral body in the intermediate and caudal series, with the exception of C15. C12 is the only vertebra that preserves both costal processes, with the other vertebrae apparently having lost these delicate structures before or after diagenesis. A single process is preserved on C11. Bourdon and Lindow (2015) report costal processes for all vertebrae in the cranial and intermediate series. The unfused carotic processes of the intermediate series are prominent in the neotype, as well as in MGUH 26770 (Bourdon and Lindow, 2015), creating a deep U-shaped carotic sulcus similar to *Eudromia*. A small and indistinct spinous process appears on C12, whereas the spinous processes of the caudal series are more robust. In C15, it is tall and block-like, although smaller than those of the thoracic series. The postzygopophyses of the caudal series are rounded and robust, and are slightly tapered in C15. The vertebral corpi of all vertebrae preserved here are bilaterally concave, but are especially so in C8/C9. Those in the caudal series are less concave than those in the intermediate series. Vertebrae in the intermediate series are longer than those of the caudal series. C10 is slightly crushed dorsoventrally. Both C14 and C15 bear a hypopophysis, which is slightly larger in C15.

### Thoracic vertebrae (Figure 4)

Leonard et al. (2005) reported nine thoracic vertebrae for MGUH 26770, whereas Bourdon and Lindow (2015) reported eight for the same specimen. Here we follow the thoracic vertebral count of Bourdon and Lindow (2015). As in other lithornithids and unlike the condition in tinamous, the thoracic vertebrae do not form a notarium (Houde, 1988, Mayr, 2008, Bourdon and Lindow, 2015). The zygapophyses are subtriangular and blunted, and reduced in comparison with those of the cervical vertebrae. Both T1 and T2 bear a hypopophysis, which is damaged in T1. The transverse processes, when preserved, are robust. The left transverse process is missing in T2 and T8. In dorsal view, these processes are widest in T1 and T2, and are directed posterolaterally as opposed to laterally in T5/T6 and T8. The spinous processes are large and subrectangular in shape, and longer dorsally than they are ventrally. They are greatest in length in T2 and T5/6. The dorsal margin of the spinous process is flat in all except T8, where it is distinctly curved. The postzygapophyses are longer and tapered in the cranial thoracic vertebrae, and rounded in the caudal thoracic vertebrae. The vertebral corpi are biconcave in all the vertebrae, but especially T2 and T5/6. It is widest and shortest in T8. We cannot clearly observe in the scan the lateral pneumatic foramina reported in the thoracic vertebrae of lithornithids (Houde, 1988, Bourdon and Lindow, 2015, Mayr, 2021b) due to infilling by matrix.

### Caudal vertebrae (Figure 5)

Two indeterminate free caudal vertebra are preserved. Houde (1988) stated that the free caudals of lithornithids “virtually lack transverse processes”, a description we agree with here. The spinal processes are low and blunted, as are the zygapophyses.

### Scapula (Figure 6, Table S1)

The right scapula of the neotype is preserved in lateral view, thus the medial view of this element is presented here for the first time. It is nearly complete, though the distinctive hooked acromion considered diagnostic of Lithornithidae is missing (Houde, 1988, Bourdon and Lindow, 2015, Mayr and Kitchener, 2025), with clear damage to the bone’s surface in that area suggesting it was broken off. Even without the distinctive hooked process, the acromion is bulbous and prominent, projecting cranially to the coracoid tubercle. We did not observe the lateral pneumatic foramen and small dorsal tubercle on the acromion reported for MGUH 26770 by Bourdon and Lindow (2015). The surface of the scapular head between the glenoid and the acromion is slightly concave. The glenoid facet is elongate and ovoid in shape, and also slightly concave. It is thicker caudally, such that the surface of the facet is angled laterally in dorsal view. The glenoid facet is continuous cranially with the coracoid tubercle, which is large and globose. A scapulotricipital tubercle is absent. There is a prominent pneumatic foramen on the medial side of the scapular head, a feature also present in tinamous (Bertelli, 2017, Widrig et al., 2023). As with other lithornithids, the body of the scapula is narrow and does not show a strong curve (Bourdon and Lindow, 2015), similar to the condition in *Eudromia*, and becomes dorsoventrally narrow distally (Nesbitt and Clarke, 2016). The left scapula is missing.

**Figure 6.**
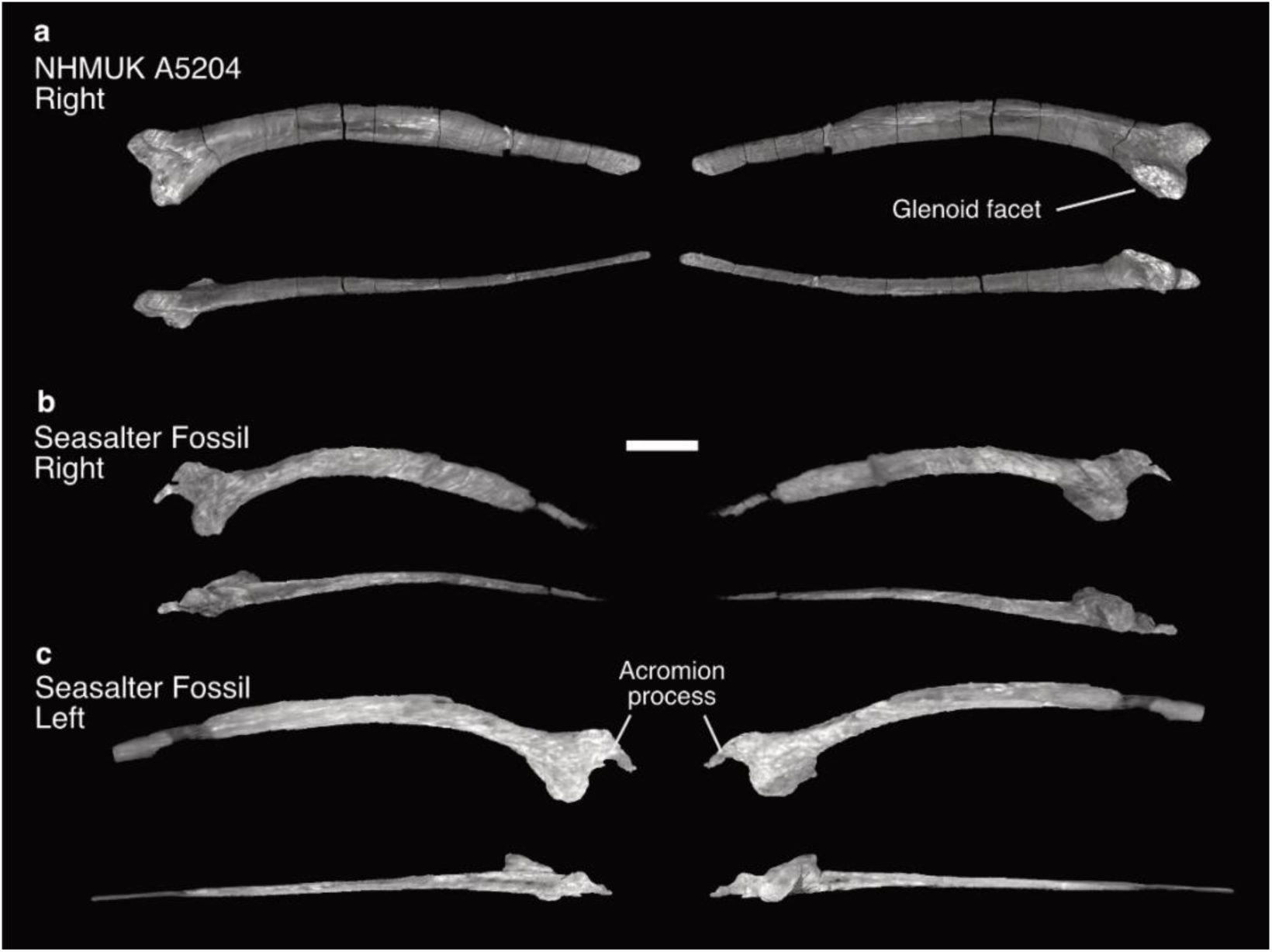
Scapulae of *Lithornis vulturinus*. (a) NHMUK A5204 right sternum, (b) Seasalter fossil right sternum, (c) Seasalter fossil left sternum; views for each in clockwise order: medial, lateral, ventral, and dorsal. Scale bar equals 10 mm.

### Humerus (Figure 7, Table S2)

The humerus is preserved in cranial view, therefore this is the first description of the humerus in caudal view. The proximal head of the humerus, the caput humeri of Baumel and Witmer (1993), is large. It is convex and ovoid in proximal view, and is positioned dorsally relative to the dorsoventral midline of the bone. Unlike the condition in *Eudromia* in which the dorsal and ventral ends of the humeral head are blunt in proximal view, in the neotype they are tapered. The cranial face of the humeral head is flat, whereas the caudal surface is convex. The area on the cranial surface between the humeral head and humeral shaft is smooth, lacking a clear coracobrachial impression. The transverse ligament sulcus is shallow, and separates the proximal bicipital crest from the cranial portion of the ventral tubercle. It does not extend dorsally past the humeral head. As remarked on by Stidham et al. (2014), the space on the caudal surface between the humeral head and shaft is also smooth, with no discernible caudal margin or “lip” distal to the humeral head. The capital incision, the incisura capitis of Baumel and Witmer (1993), forms a notch between the ventral tubercle and the humeral head in proximal view, as is the condition in other lithornithids (Houde, 1988, Stidham et al., 2014). The area just caudal to the incision is damaged, creating the appearance of a fossa that does not exist in other specimens. The deltopectoral crest is well defined and arcuate, and as in most crown birds it is curved in the dorsocranial direction (Serrano et al., 2020, Benito et al., 2022a). A concave depression extends across the distal half of the crest’s caudal surface. Between the proximal part of the deltopectoral crest and the distal part of the humeral head in cranial view is a small concavity, postulated by Bourdon and Lindow (2015) to be present across the whole of Lithornithidae. The dorsal tubercle is small and does not project far from the surface of the proximal part of the caudal humerus, in contrast to the large, ovoid ventral tubercle. The ventral tubercle is more rounded and prominent than that of *Eudromia*. The pneumotricipital fossa is large and nearly circular. It is bounded by a shallow ventral crest, but the dorsal crest is missing due to damage. The caudal margin is low and not well defined. The bicipital crest is prominent, and its ventral surface forms a U-shaped curve in both cranial and caudal view. The shallow furrow on the intumescentia between the distal portion of the bicipital crest to the dorsal margin of the transverse sulcus described by Bourdon and Lindow (2015) is present. As commented on by other authors, the humeral shaft is sigmoid in cranial and caudal view, which represents a marked departure from the relatively straight humeral shafts of tinamids (Houde, 1988, Bourdon and Lindow, 2015).The distal end of the humerus is missing, having apparently been broken.

**Figure 7.**
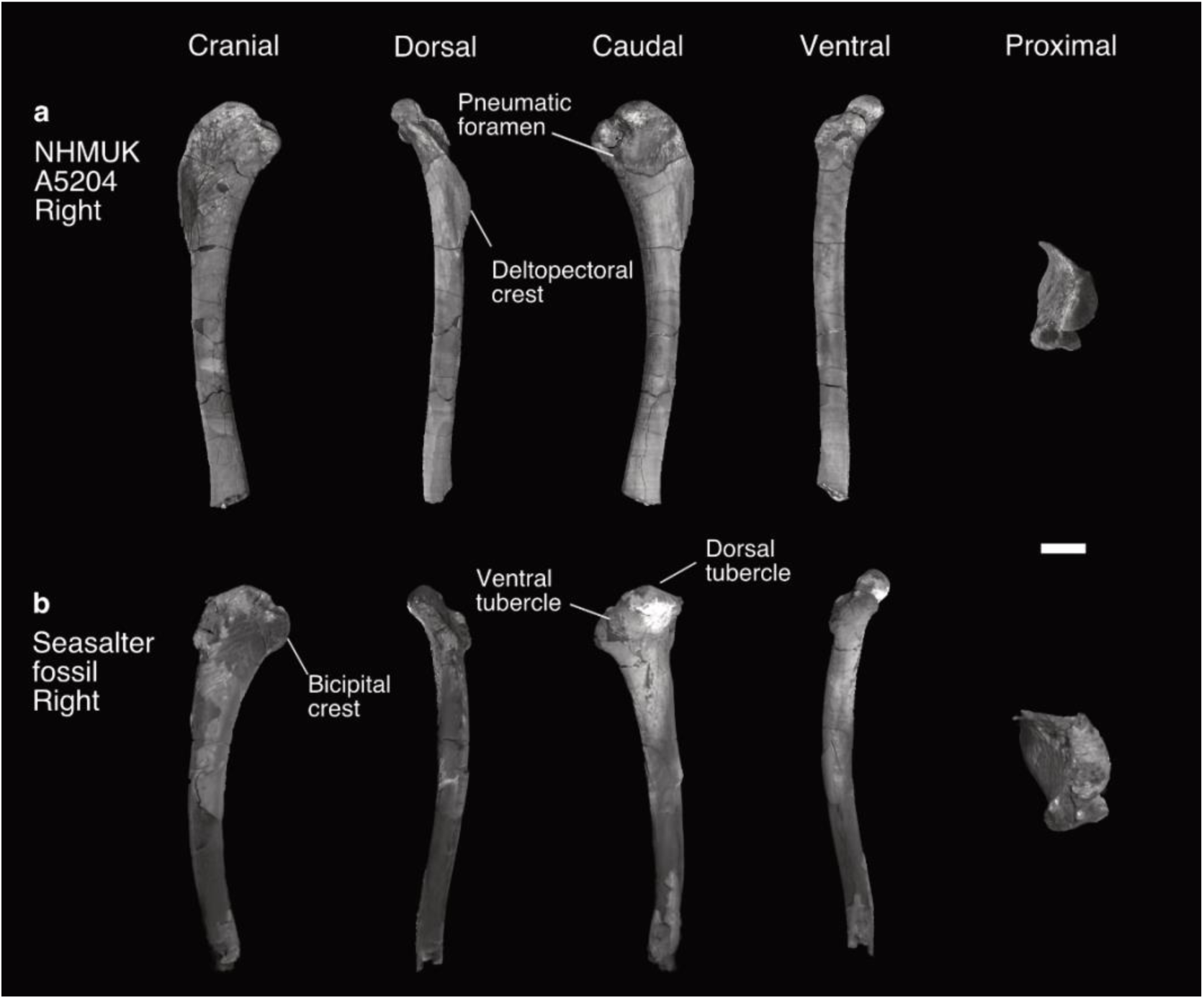
Humeri of *Lithornis vulturinus*. (a) NHMUK A5204 partial right humerus, (b) Seasalter fossil partial right humerus; in cranial, dorsal, caudal, ventral, and proximal views. Scale bar equals 10 mm.

### Radius (Figure 8, Table S3)

The right radius is preserved in cranial view missing its distal end due to breakage. The humeral cotyle is flat, and exhibits a rounded crescent shape in proximal view.

**Figure 8.**
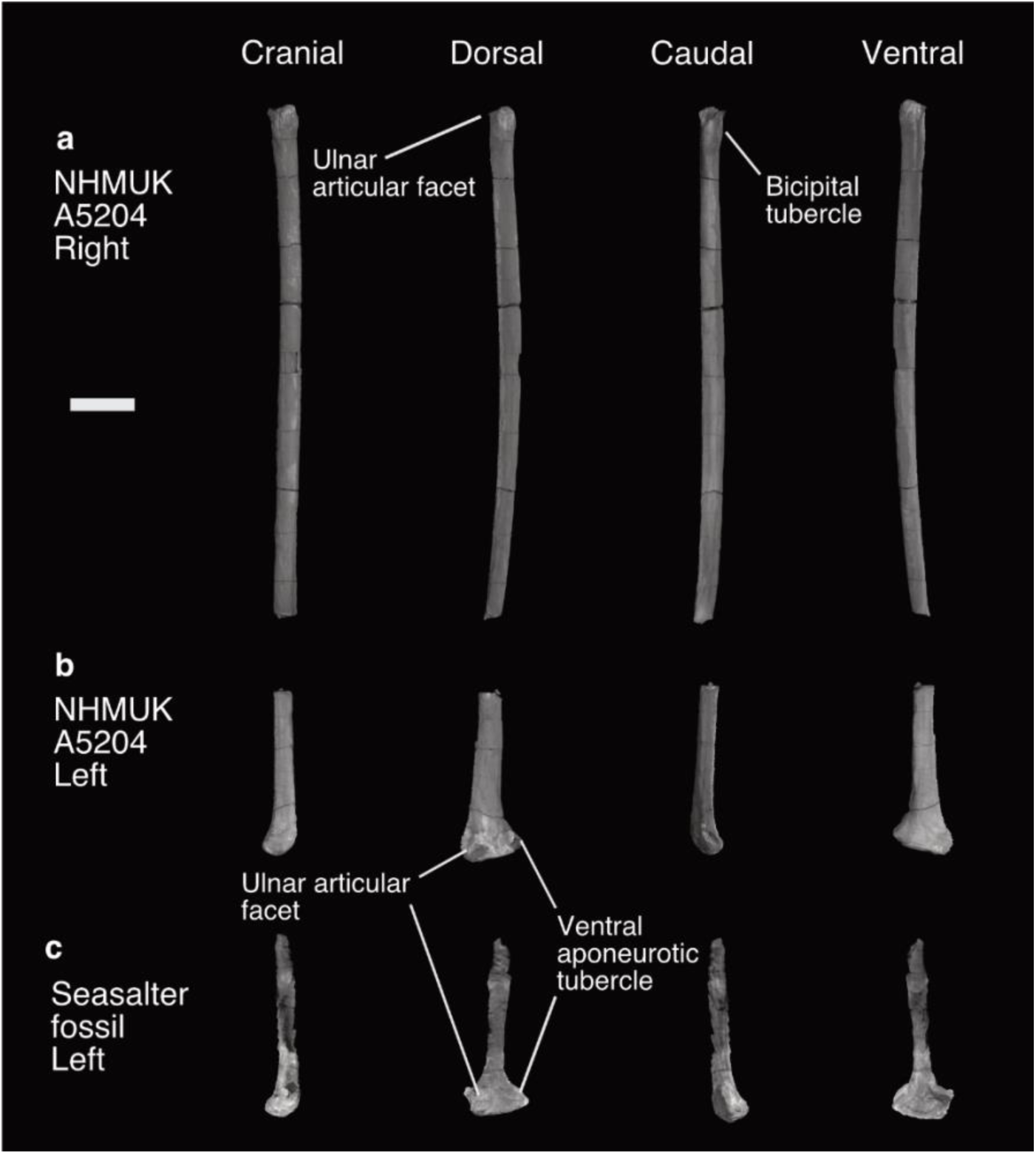
Radii of *Lithornis vulturinus*. (a) NHMUK A5204 partial right radius, (b) NHMUK A5204 distal left radius, (c) Seasalter fossil distal left radius; in cranial, dorsal, caudal, and ventral views. Scale bar equals 10 mm.

There is a small tubercle on the proximocaudal side of the ulnar articular facet. The radial bicipital tubercle is low and rounded, as in *Eudromia*. The radial shaft shows a slight curve in dorsal and ventral views. The preserved distal part of the left radius includes the distal head and a portion of the shaft. In dorsal view, the ventral aponeurotic tubercle is damaged, the tip having apparently been broken off, but still prominent. The tendinal sulcus is broad, and shallower than that of *Eudromia*. In ventral view, the ligamental depression is likewise shallow. There is no defined ulnar articular facet.

### Ulna (Figure 9, Table S4)

The proximal part of the right ulna is preserved along with the majority of the ulnar shaft. The olecranon process is low and rounded, and slightly more blunt than the condition in tinamous (Widrig et al., 2023).

**Figure 9.**
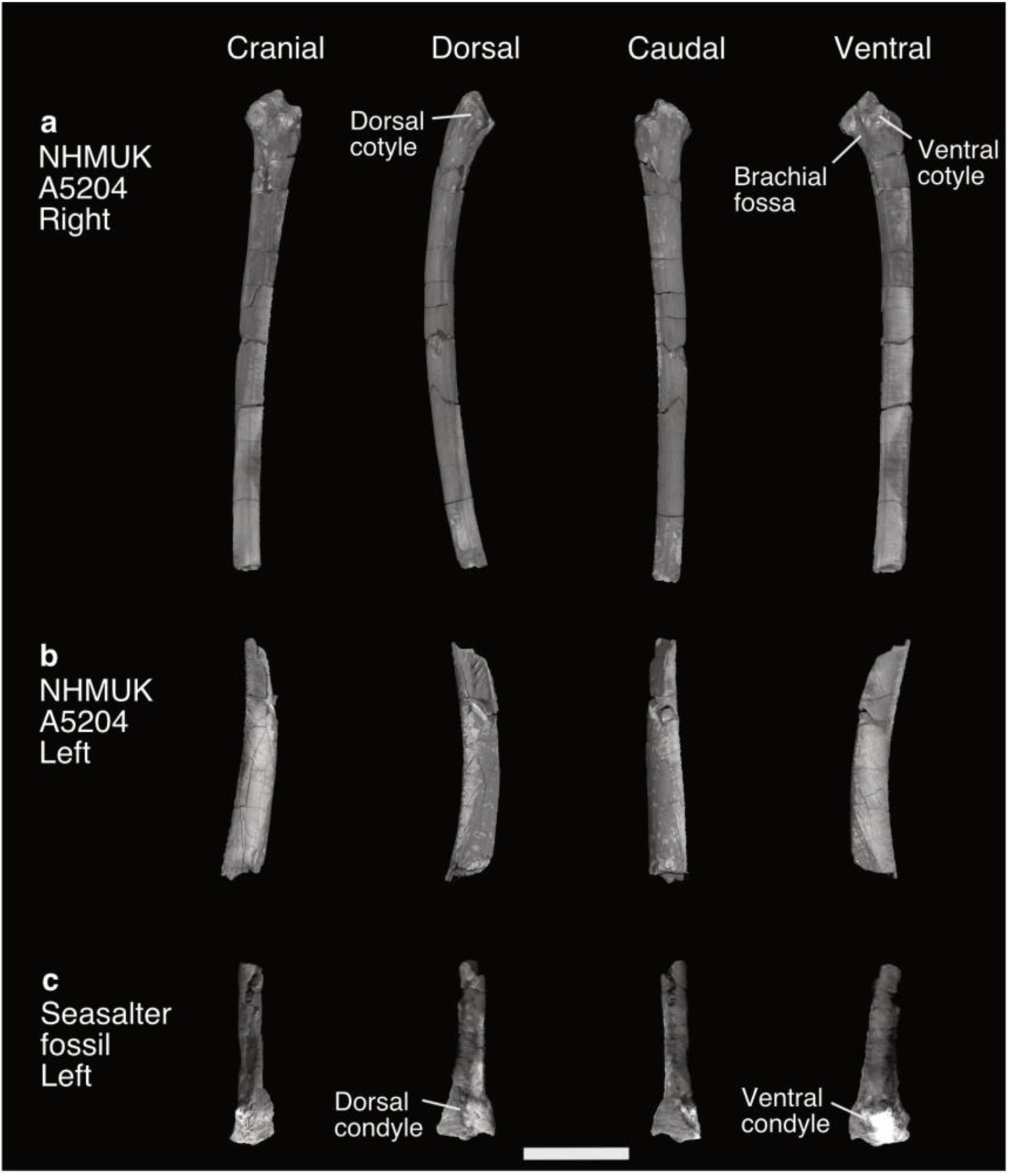
Ulnae of *Lithornis vulturinus*. (a) NHMUK A5204 partial right ulna, (b) NHMUK A5204 distal left ulna, (c) Seasalter fossil distal left ulna; in cranial, dorsal, caudal, and ventral views. Scale bar equals 20 mm.

The radial incisure is proximodistally longer and more defined than that of *Eudromia*. It is situated between the dorsal and ventral cotyles, just distal to the termination of the intercotylar crest. The dorsal cotyle is roughly triangular in proximal view. The dorsal cotylar process is straight rather than curved, and tapers to a distocranially directed point. The m. scapulotriceps impression on the dorsal surface of the dorsal cotylar process (Baumel and Witmer, 1993) is shallow with indistinct edges. The ventral cotyle is roughly quadrangular in proximal view, contrasting with the condition in *Eudromia* where it is distinctly rounded. It is approximately the same size as the dorsal cotyle. The intercotylar crest is more sharply defined than that of *Eudromia*. The brachial impression is shallow, with poorly defined edges. The collateral ventral ligament tubercle is small. The ulnar shaft is slightly curved, and almost perfectly ovoid in cross section. No remigial papillae are visible, unlike in MGUH 26770 (Bourdon and Lindow, 2015). As with the radius, the right ulna is also missing the distal end. A curved distal left ulnar shaft missing the distal head is also preserved.

### Radiale (Figure 10)

The right radiale is visible at the surface of the prepared nodule whereas the left has been fully separated from the matrix, though neither was mentioned by Houde (1988). Some surface features of the left radiale have been lost to weathering, whereas the right radiale is relatively pristine. The radiale of the *L. vulturinus* neotype does not resemble the “heart shaped” outline of this bone in cranial view in tinamids, and is instead similar to those of most neognaths and *Ichthyornis dispar* in that it is narrower in the proximodistal direction than in the dorsoventral direction, and is tapered dorsally (Mayr, 2014, Benito et al., 2022a). The dorsal tip of the right radiale is slightly abraded. The left and right radiale of the neotype are congruent with those in other members of the genus *Lithornis* (Bourdon and Lindow, 2015).

**Figure 10.**
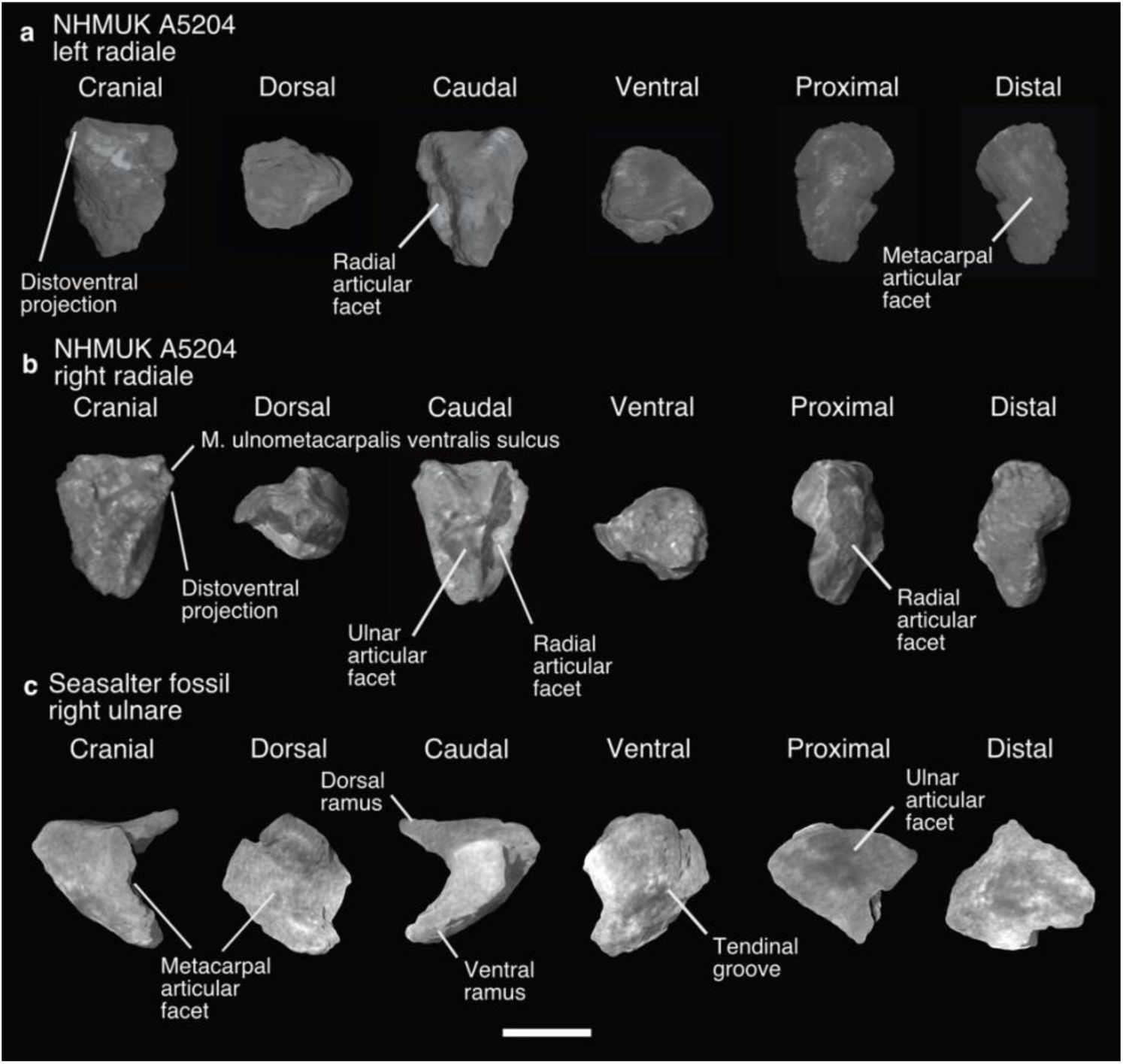
Free carpal bones of *Lithornis vulturinus*. (a) NHMUK A5204 left radiale, (b) NHMUK A5204 right radiale, (c) Seasalter fossil right ulnare; in cranial, dorsal, caudal, ventral, proximal, and distal views. Scale bar equals 5 mm.

On the distal side of the radiale, the metacarpal articular surface is rhomboidal in distal view, smooth, and slightly concave, whereas on the proximal side the radial articular surface is deeply excavated. In caudal view, the ulnar articular facet is also well excavated, even more so than the radial articular surface, though it is smaller and more rounded than the former. In cranial view, the distoventral projection is not well defined in comparison with most neognaths (Mayr, 2014), and the ulnometacarpalis ventralis muscle sulcus spans its distal margin. This position is unusual and was not observed in any of the higher level taxa examined by Mayr (2014), nor in *Ichthyornis* (Benito et al., 2022a), in which the sulcus was located on the ventral margin of the radiale. The sulcus for the extensor carpi radialis tendon is broad and shallow like that of most birds (Mayr, 2014), and crosses the centre of the radiale in the dorsoventral direction. This sulcus is not elevated from the surface of the bone, as it is in many birds examined by Mayr (2014). A ridge abuts the metacarpal articular facet.

### Carpometacarpus (Figure 11, Table S5)

Although it is visible from the external surface of the nodule, the partial right carpometacarpus was not mentioned by Houde (1988) in his description of the neotype. It is a portion of the right proximal part of the carpometacarpus missing its proximal-most portion, including the alular metacarpal and extensor process, as well as the distal parts of the major and minor metacarpals. Both the major and minor metacarpal are relatively uncurved, as is typical for lithornithids (Houde, 1988). There is no intermetacarpal process. The tuber on the proximal portion of the minor metacarpal present in all other lithornithids with the exception of *Paracathartes* (Mayr and Kitchener, 2025) is seemingly absent.

**Figure 11.**
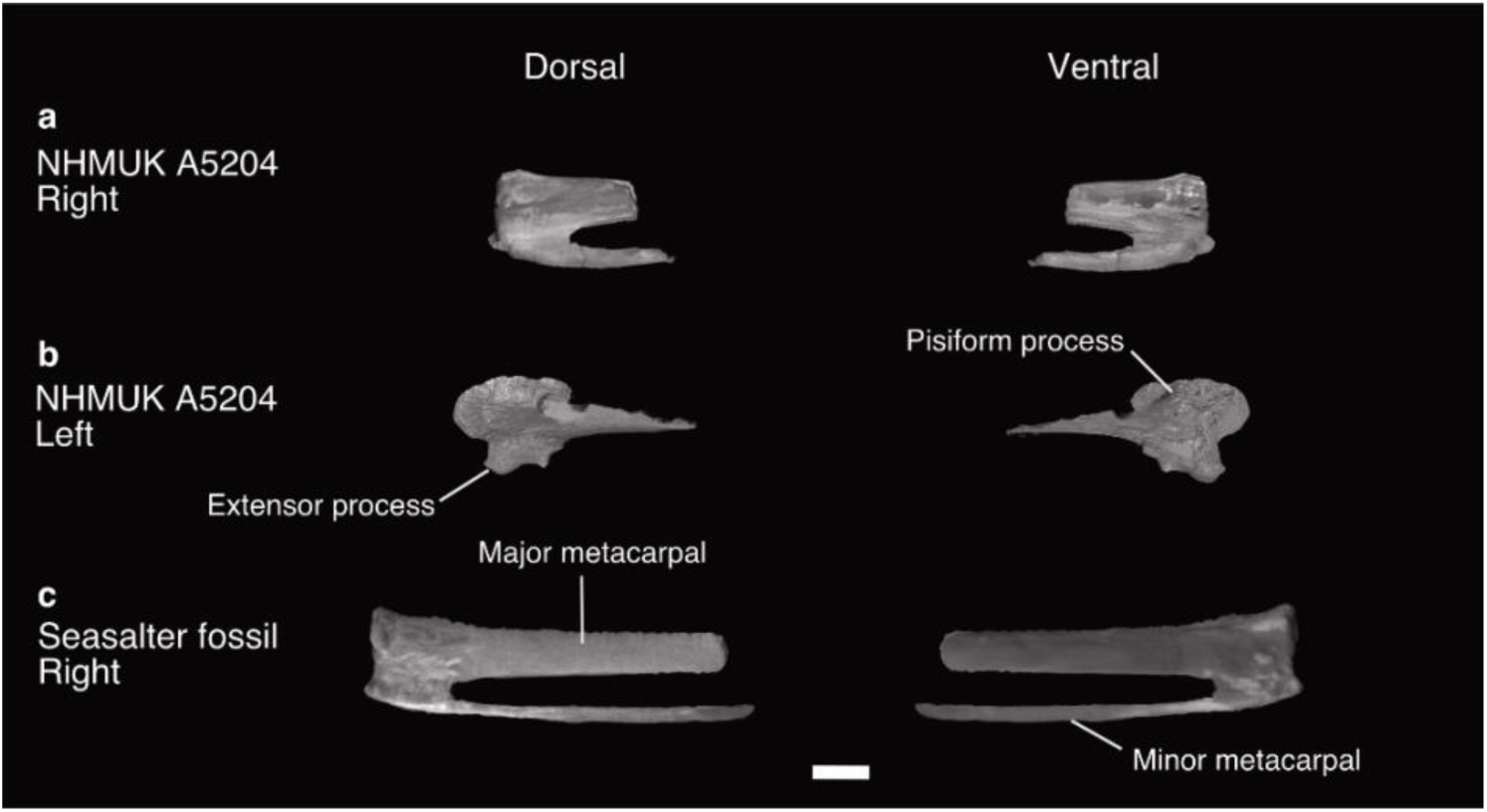
Carpometacarpi of *Lithornis vulturinus*. (a) NHMUK A5204 partial right carpometacarpus, (b) NHMUK A5204 partial left carpometacarpus, (c) Seasalter fossil partial right carpometacarpus; in dorsal and ventral views. Scale bar equals 5 mm.

A portion of the left proximal part of the carpometacarpus was scanned separately from the main nodule in a collection of small bone fragments. The proximal-most portion is preserved, as well as a portion of the major metacarpal. The ulnocarpal articular facet is missing. The extensor process is prominent, with a blunt distal end. Like all other lithornithids but unlike tinamous (Nesbitt and Clarke, 2016), the infratrochlear fossa deeply excavates the proximal surface of the pisiform process. There is a deep fossa posterodistal to the pisiform process, which again is found in all other lithornithids.

### Sternum (Figure 12, Table S6)

The right half of the sternum and the dorsal-most portion of the sternal carina are preserved. The mediolaterally broad coracoid sulci cross at the midline, with the right sulcus ventral to the left. There is no labrum tubercle at the posterior margin of the coracoid sulcus. An internal spine of the sternum is absent, whereas an external spine is present. Little can be said of the shape of the sternal carina or the presence of a supracoracoideus impression as most of the ventral portion is missing, but the carina clearly extends all the way to the caudal margin. Four intercostal incisures can be discerned caudal to the posterolaterally directed craniolateral process. The lateral margin diverges posteriorly as noted by Houde (1988), and the caudal margin would be convex had the left side of the sternum been preserved. There are no medial or lateral incisures or fenestrae, nor are there lateral trabeculae, and as such the sternum bears little resemblance to that of tinamids (Widrig et al., 2023).

**Figure 12.**
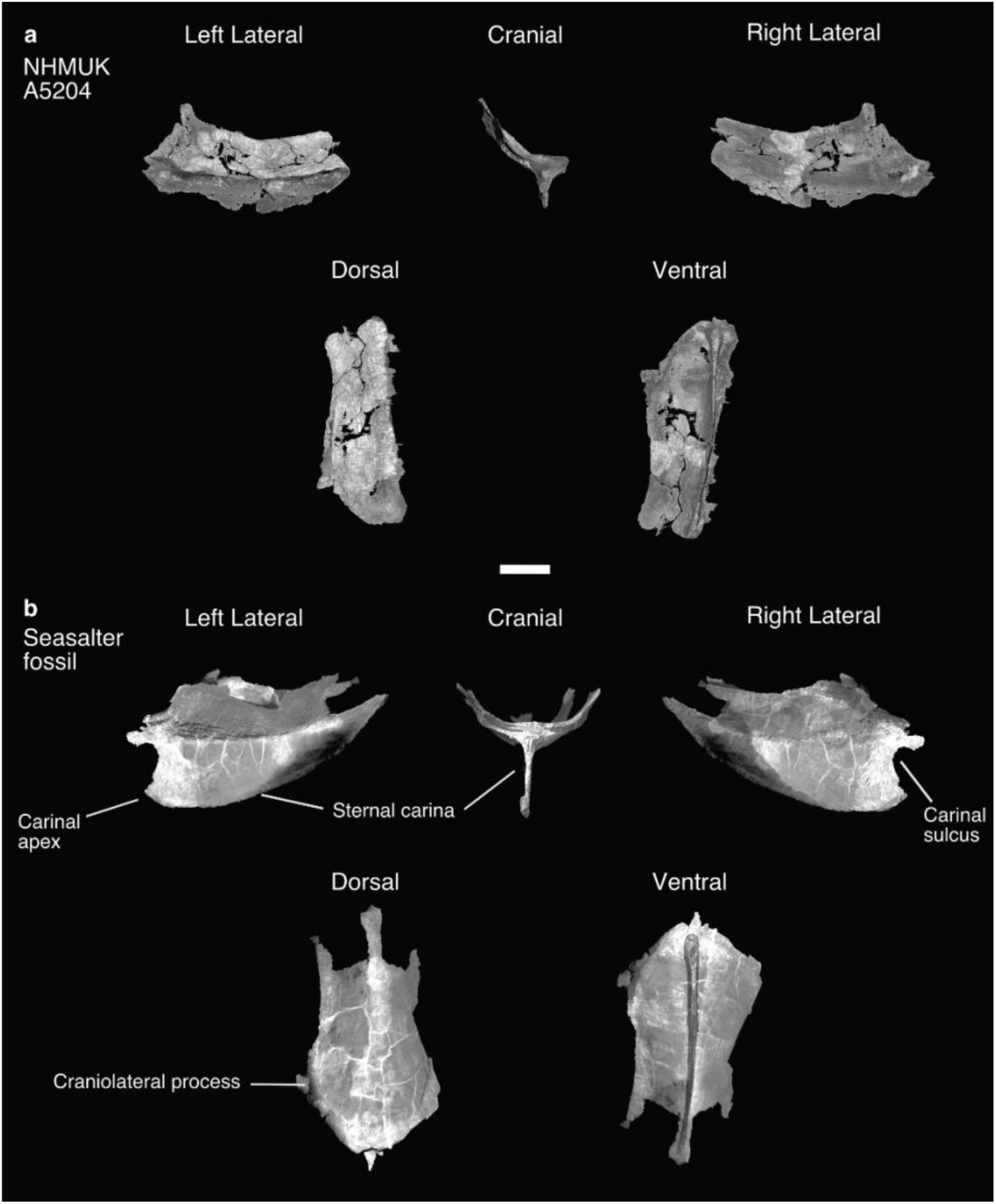
Sterna of *Lithornis vulturinus*. (a) Sternum of NHMUK A5204, (b) sternum of Seasalter fossil; in left lateral, right lateral, cranial, dorsal, and ventral views. Scale bar equals 15 mm.

### Ribs (Figure 5)

Fifteen thoracic ribs, two sternal ribs, and one isolated uncinate process were found within the nodule. None are articulated with the sternum or vertebrae. All the ribs are missing uncinate processes, which in lithornithids are not fused to the ribs (Nesbitt and Clarke, 2016). Ribs of the posterior thoracic series are broader ventrally, which was also noted for *Calciavis grandei* by Nesbitt and Clarke (2016).

### Pelvis (Figure 13)

A fragmentary right pelvis consisting of the acetabular foramen and its immediate surroundings is preserved in a separate fragment from the main nodule. Nearly all of the preacetabular portion of the ilium is missing save for a thin sliver of the medial portion of this bone. Posterior and dorsal to the nearly circular acetabular foramen is the badly damaged antitrochanter. The proximal-most portions of the right ischium and pubis are preserved in articulation. The pectineal process of the pubis is rounded and does not project far anteriorly of the contact with the ilium, similar to that of *Calciavis grandei*, *Lithornis plebius*, and *Lithornis promiscuus* (Nesbitt and Clarke, 2016) and unlike the condition seen in *Eudromia*, in which this process shows a much greater anterior projection.

**Figure 13.**
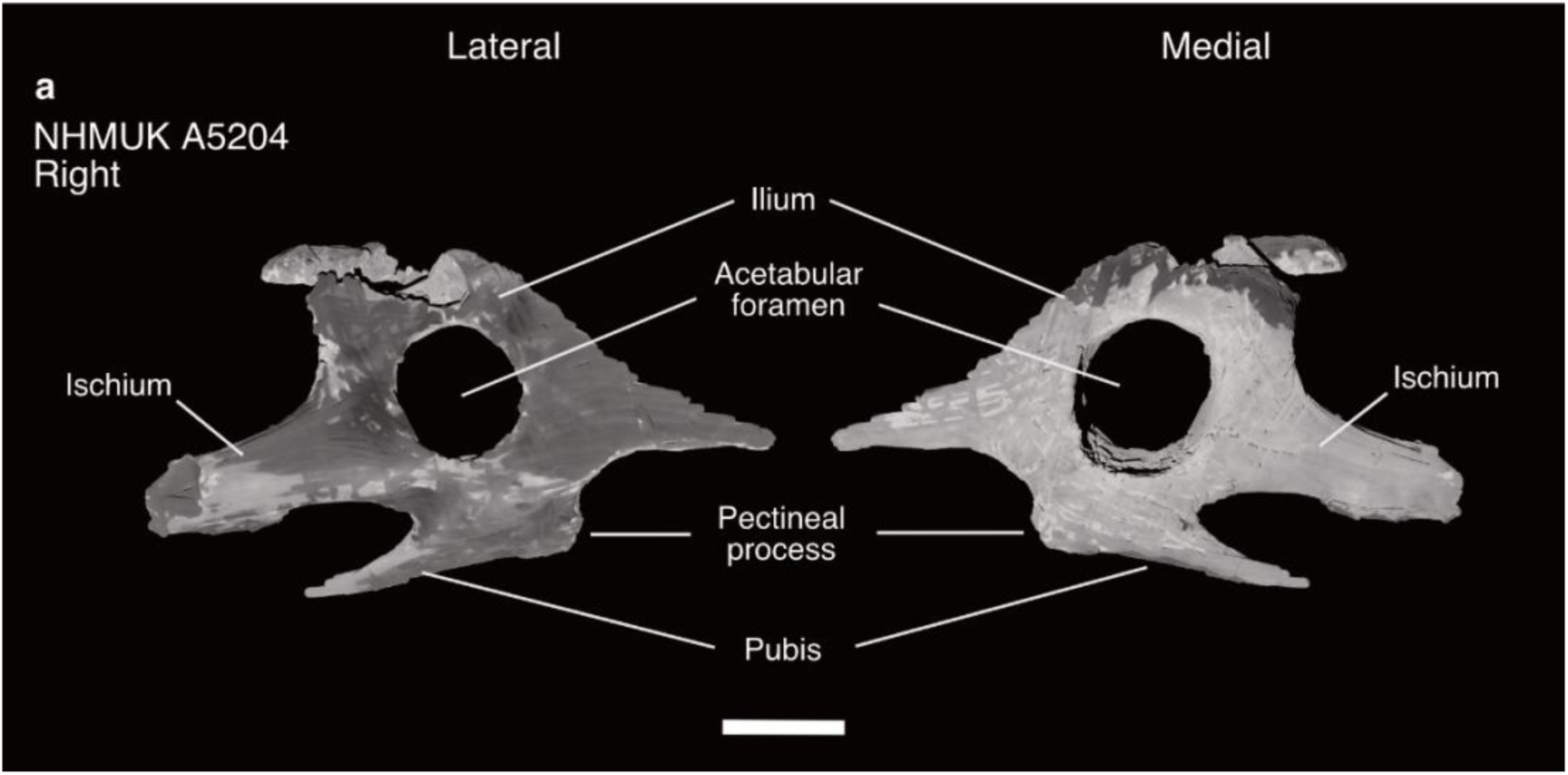
Pelvis of Lithornis vulturinus neotype NHMUK A5204. (a) Partial right pelvis; in lateral and medial views. Scale bar equals 5 mm.

### Femur (Figure 14, Table S7)

The neotype contains two incomplete femora: a weathered fragment of the proximal part of the right femur, and the distal part of the left femoral head with a portion of the shaft in slightly better condition. The neck of the rounded proximal part of the femoral head is distinct from the body of the proximal part of the femur, slightly moreso than in *Eudromia* and similar to that of *L. promiscuus*, *L. plebius*, and *C. grandei* (Houde, 1988, Nesbitt and Clarke, 2016). An oval-shaped pit on the cranial side of the femoral neck appears to represent damage to the fossil’s surface. The acetabular articular facet is slightly flattened. The trochanteric crest is much lower and less defined than that of *Eudromia*, which may be due in part to weathering, though this feature is low and not well projected in other lithornithids to begin with (Houde, 1988, Nesbitt and Clarke, 2016). The antitrochanter articular facet is shallow.

**Figure 14.**
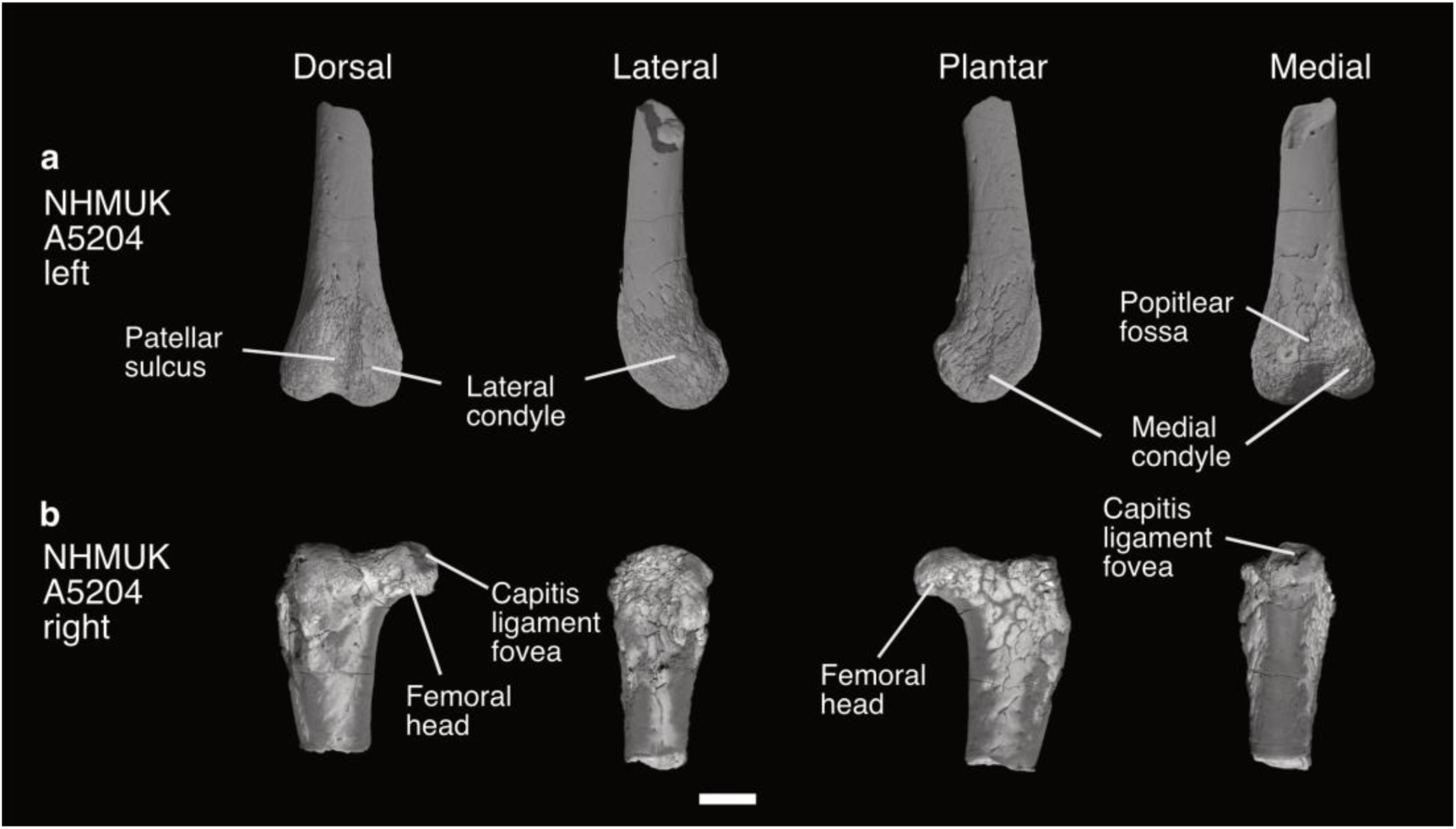
Femora of *Lithornis vulturinus* NHMUK A5204. (a) distal part of the left femur, and (b) proximal part of the right femur; in dorsal, lateral, plantar, and medial views. Scale bar equals 5 mm.

In dorsal view, the popliteal fossa of the distal part of the femur is shallow, as it is in other members of the clade (Nesbitt and Clarke, 2016). The lateral collateral ligament impression and m. tibiolis cranialis tendon fovea are both visible despite weathering to the surface of the distal femoral head. However, the lateral and medial epicondyles appear to have been removed by weathering. In plantar view, the medial supracondylar crest is visible, although it is damaged. The cruciati caudalis ligament impression is located slightly median of centre on the lateral condyle. Features such as the tibiofibular crest and the M. gastroc. lateralis tubercle are missing due to surface weathering.

### Tibiotarsus (Figure 15, Table S8)

The left proximal head of the tibiotarsus is broken proximally to the assumed positions of the fibular crest and the collateral medial ligament impression, and no other portion of the left tibiotarsus is preserved. As in all lithornithids, the cranial cnemial crest is longer than the lateral cnemial crest, whereas the lateral crest is more strongly projected than the cranial cnemial crest (Nesbitt and Clarke, 2016). The lateral crest is prominent in dorsal and plantar view and has a slight hooked shape as compared with that of *Paracathartes*, *Pseudocrypturus*, and *L. plebius*, in which the edge is rounded. It is similar in shape to that of *Calciavis grandei* and *L. promiscuus*, though this hook-like shape is slightly more pronounced in the latter. This feature unfortunately cannot be compared to MGUH 26770, as all hindlimb elements are missing from this fossil. In proximal view, the intercnemial sulcus forms a wide U shape that opens in the craniolateral direction. The patellar crest is prominent. In plantar view, the lateral articular surface is flat, whereas the fibular caput itself is prominent. The popliteal tubercle is much less defined and projected than in *Eudromia*.

**Figure 15.**
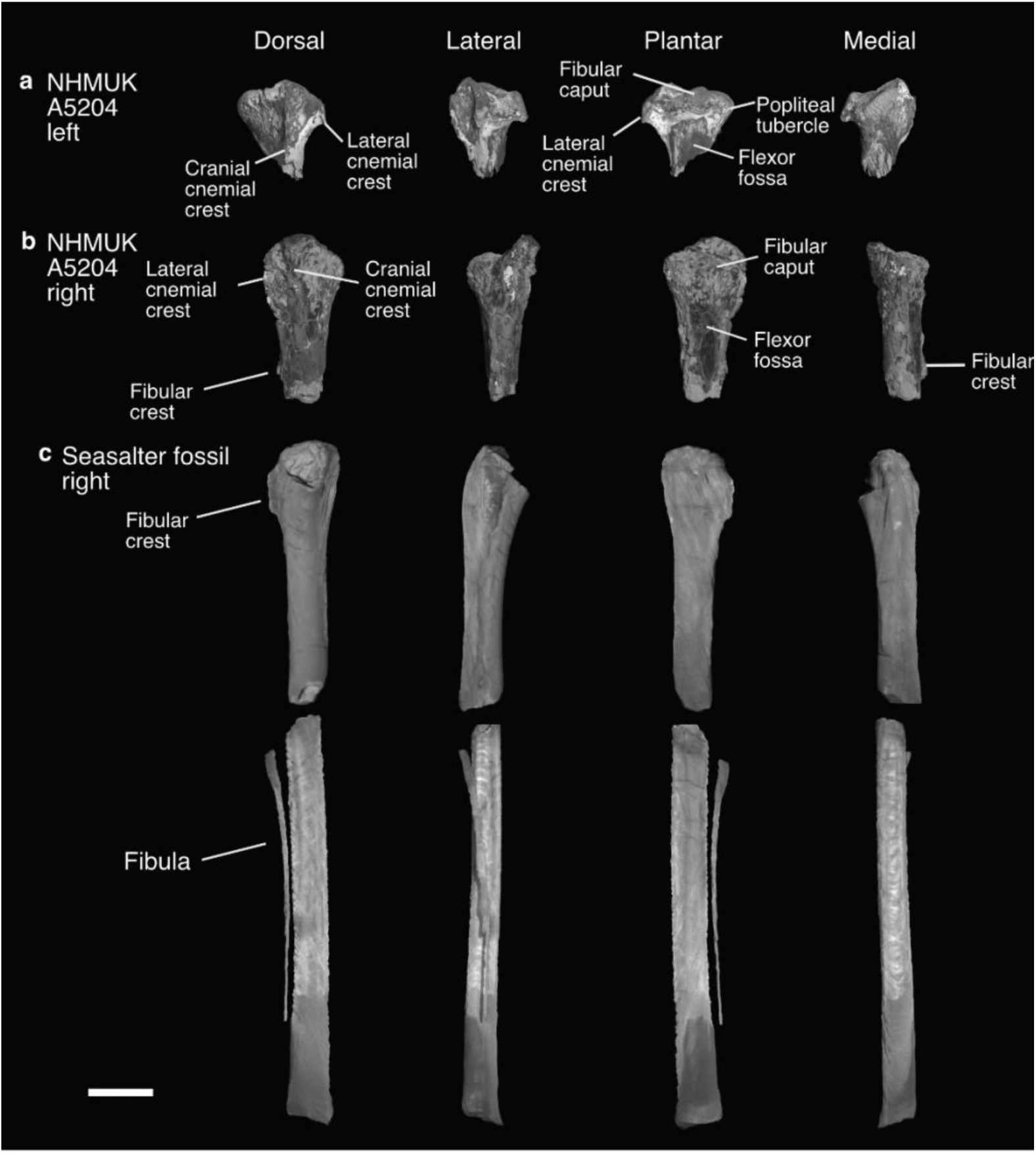
Tibiotarsi and fibula of *Lithornis vulturinus*. (a) NHMUK A5204 proximal part of the left tibiotarsus, (b) NHMUK A5204 proximal part of the right tibiotarsus, (c) Seasalter fossil proximal part of the right tibiotarsus, tibiotarsus shaft, and fibula; in dorsal, lateral, plantar, and medial views. Scale bar equals 10 mm.

Unlike the left tibiotarsus, a small portion of the shaft is preserved along with the proximal head of the right tibiotarsus, but many surface features have been damaged by erosion or lost entirely. Nearly all of the fibular crest is missing save for a small area near the distal end of the bone. The lateral cnemial crest has been rounded off by weathering.

### Pedal phalanges

Only a single pedal phalanx was found, with the proximal end broken. The pit on one side of the phalanx for the collateral ligaments is much deeper than that on the other, although medial and lateral directions are challenging to ascertain; the missing proximal articular surface prevents us from identifying the position of this phalanx.

### Anatomical Description: New fossil (Seasalter specimen)

#### Vertebral column (Figures 5, 16, 17, 18)

A total of nineteen vertebrae are preserved. The vertebrae here agree in morphology with those of the neotype, and many are preserved in articulation.

**Figure 16.**
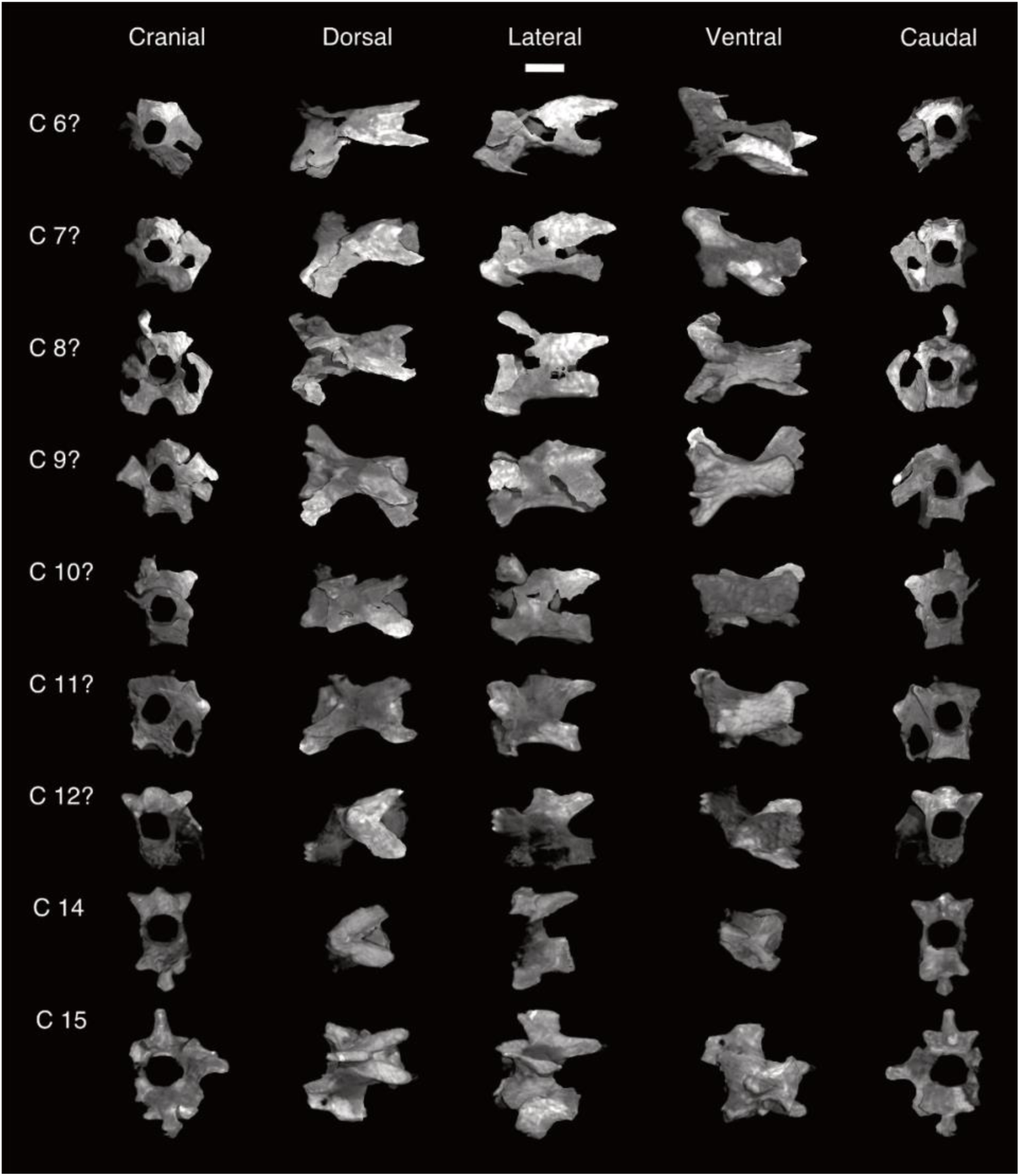
Cervical vertebrae of the Seasalter fossil. Possible 6^th^ through 12^th^ cervical vertebrae, cervical vertebrae 14 and 15; in cranial, dorsal, lateral, ventral, and caudal views. Scale bar equals 5 mm.

**Figure 17.**
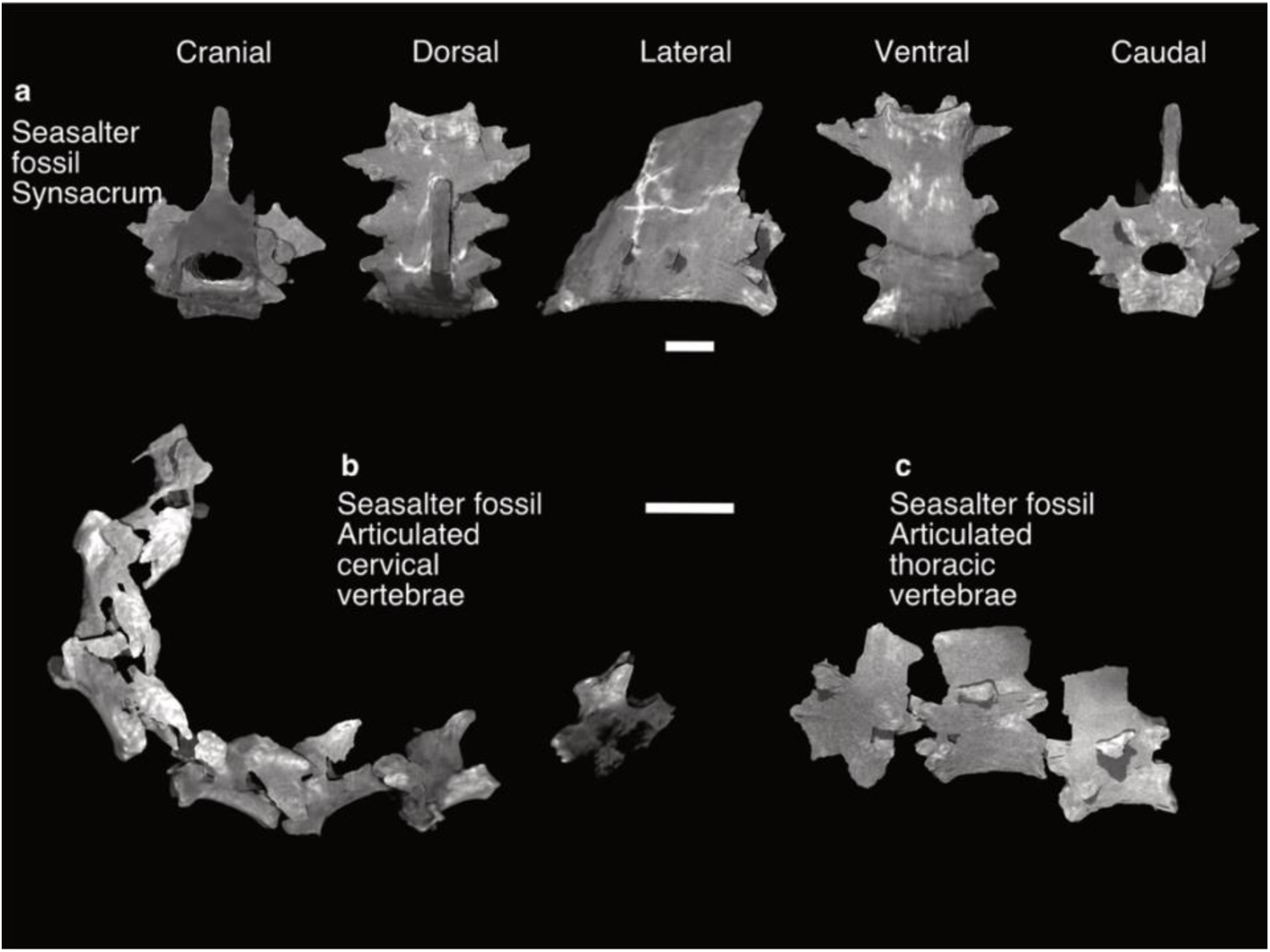
Articulated vertebrae of the Seasalter fossil. (a) Partial synsacrum in cranial, dorsal, lateral, ventral, and caudal views. Scale bar equals 5 mm. (b) Cervical vertebrae and (c) thoracic vertebrae found in articulation, in left lateral view. Scale bar equals 10 mm.

**Figure 18.**
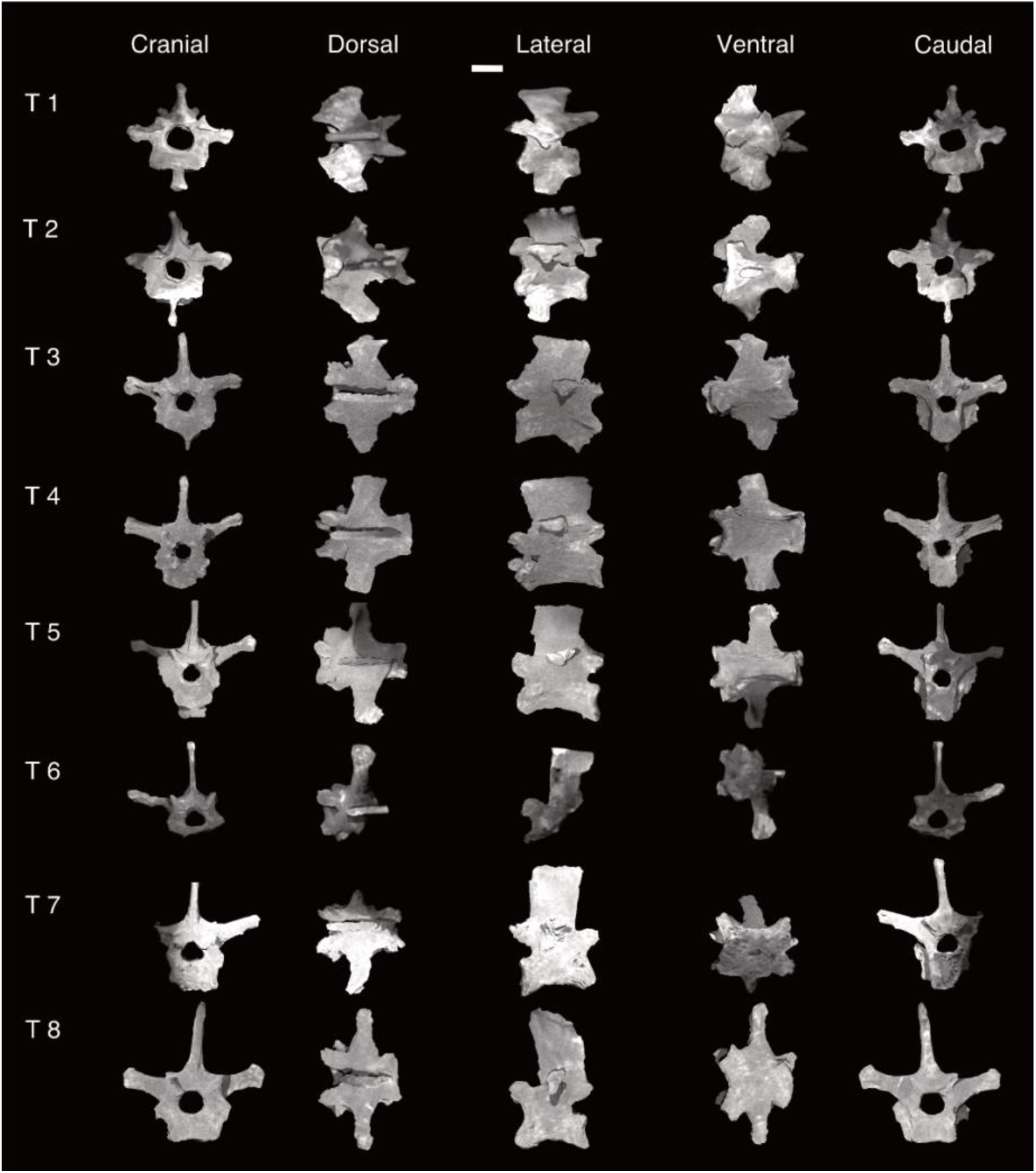
Thoracic vertebrae of the Seasalter fossil. 1^st^ through 8^th^ thoracic vertebrae; in cranial, dorsal, lateral, ventral, and caudal views. Scale bar equals 5 mm.

### Cervical vertebrae (Figures 16, 17)

Vertebrae that we have tentatively identified as C6 through C12 are preserved in articulation. C14 and C15 were found in articulation separate from the other cervical vertebrae. The cervical vertebrae are generally less well preserved than those of the neotype. C6 and C7 are the only vertebrae that preserve costal processes; these delicate structures were damaged and lost on all other cervical vertebrae preserved here. C11 bears what appears to be a very small and damaged spinous process. This process becomes larger and more distinct in C12 and C14, and in C15 it is large and subrectangular, with a curved anterior margin. C14 and C15 both bear a hypopophysis.

### Thoracic vertebrae (Figures 17, 18)

A complete disarticulated series of eight thoracic vertebra are preserved. Due to the infilling of material of the same density as the fossils themselves, the pneumatic lateral openings described by Mayr (2021b), reported as pneumatic foramina by Houde (1988), are not visible in the scans, but are visible when the fossil is observed by the naked eye, as in MGUH 26770 (Bourdon and Lindow, 2015). As expected, they do not form a notarium. The transverse processes are large, with club-like lateral ends. From dorsal view they appear posterolaterally directed in T1 and T2, and laterally directed in all others. Bourdon and Lindow (2015) observed that the transverse processes of T7 and T8 are narrower than those of the more cranial thoracic vertebrae in MGUH 26770, which we also find here. The spinous processes are tall and subrectangular, as in the neotype. They reach their greatest craniocaudal length in T4, and their greatest dorsoventral height in T7 and T8. Both the anterior and posterior margins of the spinous process of T8 are curved.

### Synsacrum (Figure 17)

The caudalmost three vertebra of the synsacrum are preserved. The cranial face of the first vertebra in this partial synsacrum has been damaged by erosion. Three pairs of transverse processes are preserved, though the individual vertebrae cannot be distinguished. The posterior-most pair of transverse processes are the largest, and all three pairs are posteriolaterally directed, though less posteriorly directed than those of *Calciavis grandei* (Nesbitt and Clarke, 2016). As in MGUH 26770 *L. vulturinus*, the sacral spinous crest is prominent. It is rhomboidal in shape, and comes to a posterodorsally directed point at its distal end. There are no distinguishing features on the ventral surface of this partial synsacrum.

### Caudal vertebrae (Figure 5)

Two indeterminate caudal vertebrae are preserved. On the basis of their larger relative size, they are likely more cranial than the caudal vertebra preserved in the neotype.

### Scapula (Figure 6, Table S1)

Both scapulae are preserved in this specimen. The left scapula is nearly complete, whereas the right is missing its distal end. The distinctive hooked acromion of lithornithids that is missing in the neotype is preserved in both scapulae. The long, narrow scapula is similar in overall shape to *Lithornis plebius*.

The surfaces of the glenoid facets are flat, with a raised tubercle on the distal ends of the facets. The distal end of the glenoid facet of both scapulae is slightly pointed in comparison to the neotype, forming a rounded teardrop shape rather than an oval.

Based upon our own observations of lithornithids in the USNM collections, we interpret this as normal intraspecific variation. The pneumatic foramen on the medial part of the scapular head can be observed on the right scapula, but not the left.

Whether this is due to a genuine absence or infilling by sediment of the same radiodensity as the fossil material is unclear. The scapular body of both is more strongly curved than that of the neotype, which is notable as Houde (1988) states that the scapular body is broader and more curved in larger lithornithids.

### Coracoid (Figure 19, Table S9)

This specimen preserves nearly complete right and left coracoids. Although preservation of the sternal margin of the left coracoid is poor due to incomplete mineralisation, it is clear that this margin was distinctively wide as in other lithornithids (Houde, 1988, Bourdon and Lindow, 2015). The right coracoid is missing the medial end of the sternal facet altogether, which appears to be the result of a combination of incomplete mineralisation and poor contrast between fossil and matrix. The scapular cotyle is deeply excavated and circular, in contrast to the shallow scapular articular surface of tinamids (Bertelli et al., 2014, Bertelli, 2017, Widrig et al., 2023, Mayr and Kitchener, 2025). The glenoid facet is cranial to the scapular cotyle, as in most crown birds (Benito et al., 2022a), and the surface of the glenoid itself is flat and rhomboidal in shape. The flange-like clavicular articular facet is present on the left coracoid, but is damaged on the right coracoid. The distal ends of the procoracoid and acrocoracoid processes are broken on the right coracoid, whereas the small procoracoid process on the left coracoid is intact. As in other lithornithids, the coracoidal body is narrow (Houde, 1988, Leonard et al., 2005, Bourdon and Lindow, 2015, Nesbitt and Clarke, 2016). A supracoracoideal nerve foramen is not visible on either coracoid, though its absence here may be due to matrix infill and poor mineralization as it is recorded in all other lithornithids with preserved coracoids (Houde, 1988, Bourdon and Lindow, 2015, Mayr and Kitchener, 2025). The supracoracoideal muscle impression is deep, but its edges are poorly defined on each coracoid. As in MGUH 26770, the lateral process is projected proximally (Bourdon and Lindow, 2015) on the left coracoid, but is missing from the right.

**Figure 19.**
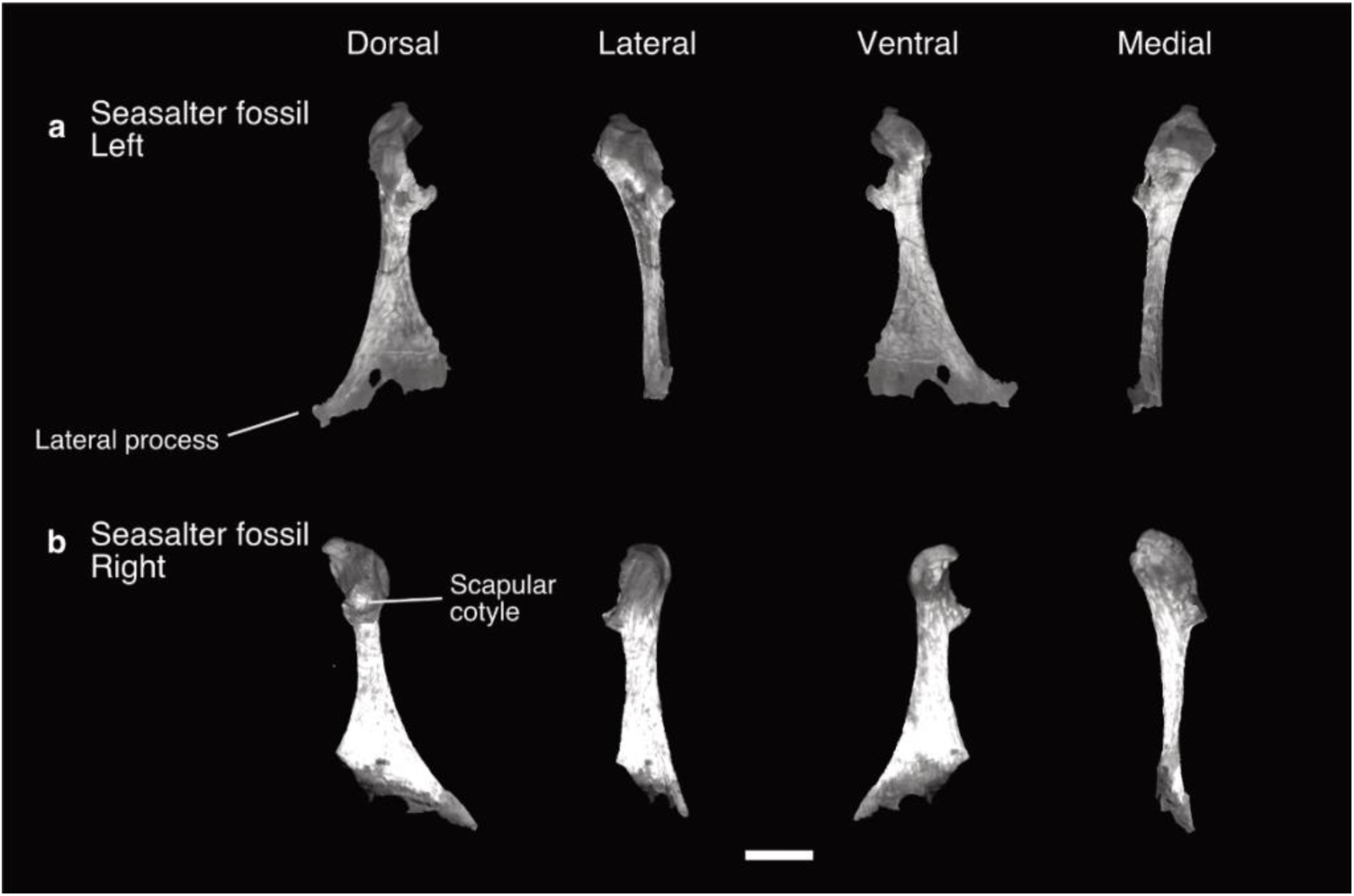
Coracoids of the Seasalter fossil. (a) left coracoid, (b) right coracoid; in dorsal, lateral, ventral, and medial views. Scale bar equals 10 mm.

### Humerus (Figure 7, Table S2)

The proximal part and shaft of the right humerus preserved and is similar in nearly all regards to that of the neotype. The shape and position of the humeral head agrees with that of the neotype. We cannot observe a coracobrachial impression, which in *Eudromia* is very shallow and indistinct. A transverse ligament sulcus is present between the proximal bicipital crest and the ventral tubercle. The area on the caudal surface between the humeral head and shaft is smooth. The dorsal tubercle is small, whereas the ovoid ventral tubercle is large and robust. Like in the neotype, the capital incision forms a notch between the ventral tubercle and the humeral head in proximal view. Due to infilling of material of the same radiodensity as the fossil itself, the pneumatic foramen is indistinct.

However, a sulcus can be observed ventrodistal to the ventral tubercle in the same location as the pneumatic foramen on the neotype. The bicipital crest is prominent and hook shaped in cranial view. The furrow on the intumescentia seen in the neotype appears to be present here, but is less distinct. The majority of the deltopectoral crest is missing, but what remains of the ventral portion agrees in shape with the arcuate crest of the neotype. There is a small concavity between the proximal part of the deltopectoral crest and the distal part of the humeral head. The only prominent difference in shape between this humerus and the neotype is the more extreme sigmoid curve of the shaft in this specimen. The distal end of the humerus is missing.

### Radius (Figure 8, Table S3)

The distal part of the left radius is preserved, albeit poorly due to incomplete mineralisation and poor contrast. The shaft is poorly preserved such that its diameter cannot be assessed. From dorsal view, the ventral aponeurotic tubercle is distinct, and the tendinal sulcus is shallow. In ventral view, the ligamental depression is also shallow.

### Ulna (Figure 9, Table S4)

Little can be said of the morphology of this distal part of the right ulna, as it is very poorly preserved. In dorsal view, the dorsal and ventral ulnar condyles can be discerned, but the tendinal incisure and carpal tubercle cannot. In ventral view, a shallow carpal tubercle incisure is visible, but the radial depression and intercondylar sulcus appear to have been lost during diagenesis.

### Ulnare (Figure 10)

A single right ulnare was the only free carpal bone found in this nodule. Overall it is in good condition, with some damage to the cranial side of the dorsal ramus (*crus breve* of Baumel and Witmer). The dorsal ramus is approximately the same length as the ventral ramus, though more slender than the latter. The protuberance situated on the ventrodistal surface of the dorsal ramus in *Lithornis promiscuus* is much reduced here, though whether this is due to weathering or a genuine reduction in size cannot be determined. The overall shape of the ulnare is similar to that of *Lithornis plebius*, which also apparently lacks this protuberance. The ventral ramus (*crus longum* of Baumel and Witmer) is robust. Both rami are gently curved in shape. A tuber is present on the caudal surface where the two rami meet, as in Galloanserae, *Lithornis promiscuus*, *Lithornis plebius*, and *Calciavis grandei* (Nesbitt and Clarke, 2016). The ulnar articular facet located on the proximal surface is slightly concave and ovoid in overall shape. The metacarpal incisure is deep and well defined. On the ventral surface, the tendinal groove is mostly obscured by weathering.

### Carpometacarpus (Figure 11, Table S5)

More of the distal part of the carpometacarpus is preserved here than in the neotype, though it is also missing the alular metacarpal, the ulnocarpal articular face, and the carpal trochlea, as well as its distal end. The right carpometacarpus of this specimen is very similar to the neotype, with the typical uncurved major and minor metacarpi of lithornithids. The pisiform process is not as prominent as that of *Eudromia*. There does not appear to be a tendinal sulcus on the major metacarpus, though this may be due to preservation as it is present on the carpometacarpi of *Lithornis celetius, Lithornis promiscuus, Pseudocrypturus cercanaxius, ?Pseudocrypturus gracilipes, ?Pseudocrypturus danielsi,* and *Paracathartes howardae* (Houde, 1988, Mayr and Kitchener, 2025). An intermetacarpal process is absent. The intermetacarpal space is elongate, narrow, and rectangular. The tuber on the proximal part of the minor metacarpal present in other members of the genus *Lithornis* is seemingly absent here.

### Sternum (Figure 12, Table S6)

With the exception of its caudal margin, costal processes, and left craniolateral process, the sternum is completely preserved and appears to be minimally distorted. The crossed coracoidal sulci are clearly visible. The prominent external spine is larger than that of the neotype. The right craniolateral process resembles the neotype, whereas the left is indistinct, likely due to poor contrast with the matrix. There appear to be four intercostal incisures on the right side of the sternum. The left side is unfortunately either damaged, poorly mineralised, or shows poor contrast in the area they would be expected to be found. The carinal apex does not project further in the cranial direction than the sternal rostrum, and the carinal sulcus is deep and well defined. The sternal keel is deepest cranially, and extends to the caudalmost end of the sternum. The “small but distinct” impression of m. supracoracoideus mentioned by Houde (1988) cannot be discerned here, as the relatively poor contrast between fossil and matrix inhibits the identification of finely detailed surface features. We cannot be certain as to the shape of the caudal margin in this fossil, as the poor mineralization of the posterior margin of the sternum has led to the appearance of deep notches contra the family and order diagnosis of Houde (1988). A portion of the posterior margin is visible on the surface of the nodule containing the fossil, and to the naked eye its appearance supports the interpretation that the apparent notches are due to a combination of incomplete mineralisation and surface weathering, rather than being representative of the original shape of the sternum.

### Ribs (Figure 5)

Twelve thoracic ribs and two sternal ribs are preserved. The uncinate processes are missing from all of the ribs but one. As in *Calciavis grandei*, ribs of the posterior thoracic series are broader ventrally (Nesbitt and Clarke, 2016).

### Tibiotarsus (Figure 15, Table S8)

A damaged proximal part of the right tibiotarsus together with the tibiotarsus shaft missing its distal end are preserved as two matching fragments. The proximal articular surface of the tibiotarsus is missing, as are the cranial and lateral cnemial crests and flexor fossa due to breakage of the proximal head. A portion of the fibular crest is preserved.

### Fibula (Figure 15)

The fibula is preserved close to its life position adjacent to the tibiotarsus shaft. The iliofibular tubercle is missing. It is present in *Calciavis grandei* as a thin phalange (Nesbitt and Clarke, 2016) and we also observed this tubercle in *Lithornis promiscuus*, *L. celetius*, *L. plebius*, *Pseudocrypturus cercanaxius*, and *Paracathartes howardae*; therefore we conclude that its absence here is due to damage.

### Tarsometatarsus (Figure 20, Table S10)

The proximal portions of the right and left tarsometatarsi are preserved. The anterior surface of the right tarsometatarsus appears somewhat worn, as it lacks the lip on the posterolateral margin of the lateral cotyle identified by Houde (1988) as a shared feature of all palaeognaths. The medial cotyle is more strongly concave than the lateral cotyle. In dorsal view, the intercotylar eminence is prominent and spherical here and in all other lithornithids, as noted by Houde (1988). The extensor sulcus is deep and lacks an m. tibialis cranialis tuberosity.

**Figure 20.**
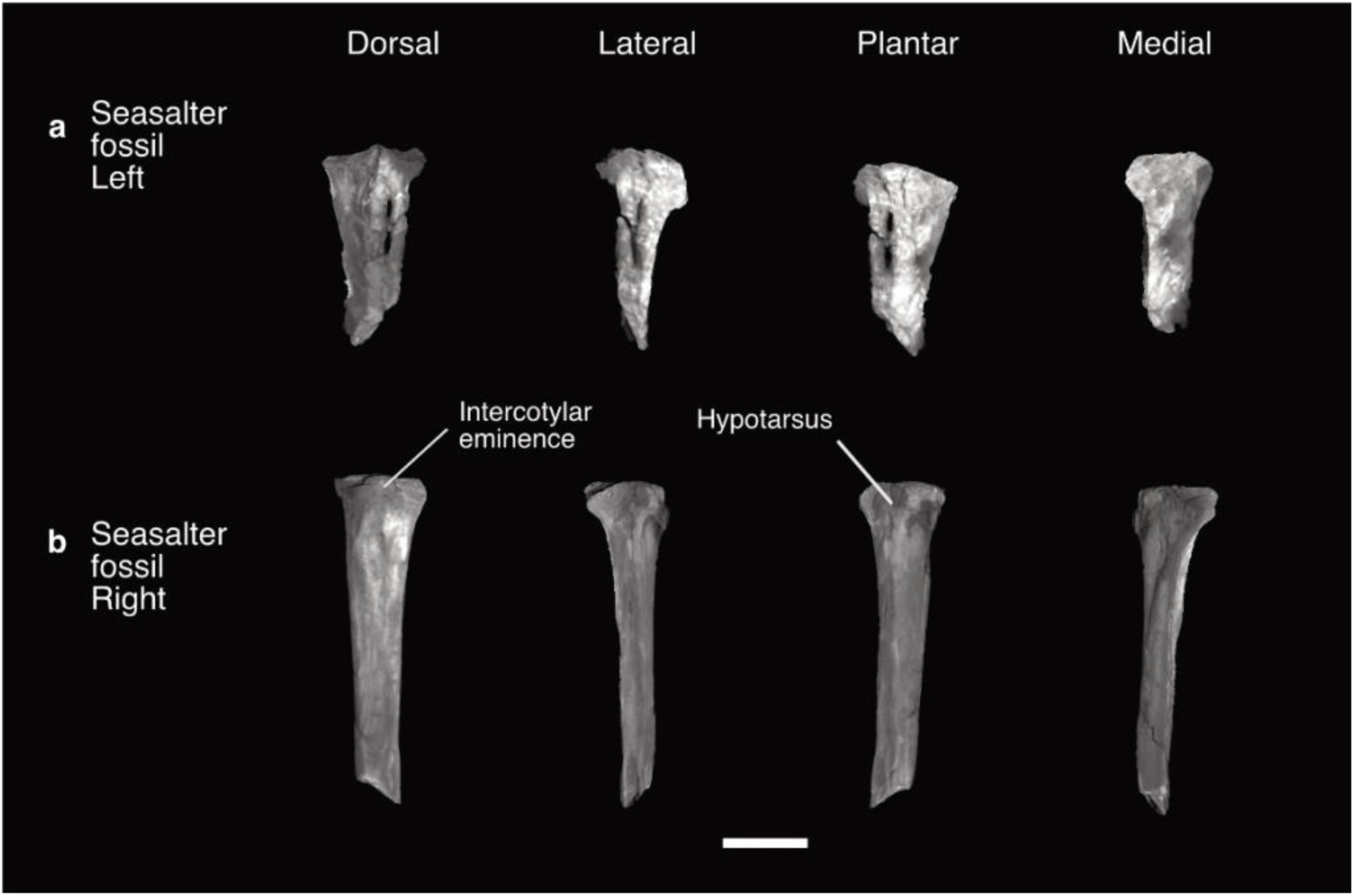
Tarsometatarsi of the Seasalter specimen. (a) Seasalter proximal part of the left tarsometatarsus, (b) Seasalter proximal part of the right tarsometatarsus; in dorsal, lateral, plantar, and medial views. Scale bar equals 10 mm.

On the plantar surface, the hypotarsus is block-like. As expected for a lithornithid (Houde, 1988, Nesbitt and Clarke, 2016, Mayr and Kitchener, 2025) no hypotarsal canals are present. The hypotarsus can be described as asulcate, as is the condition in Ichthyornithes and extant Cathartidae, Sagittariidae, and Cariamiformes (Mayr, 2016, Benito et al., 2022a). The left tarsometatarsus is in poor condition, as it was exposed on the exterior of the clay nodule. The infracotylar dorsal fossa is eroded so completely as to create the appearance of a foramen. It is otherwise consistent with the morphology of the right tarsometatarsus.

### Body mass estimates

Using the equations of Field et al. (2013), we estimated the body mass of the neotype and the new fossil from Seasalter, as well as published measurements for MGUH 26770, SGPIMH MEV1, and the NMS lithornithid collections, and from our own measurements of all lithornithids in the collections of the USNM and NHMUK with sufficient material (Figure 21). In total our sample includes estimates for 25 individuals representing eleven species (Supplementary Table S11).

**Figure 21.**
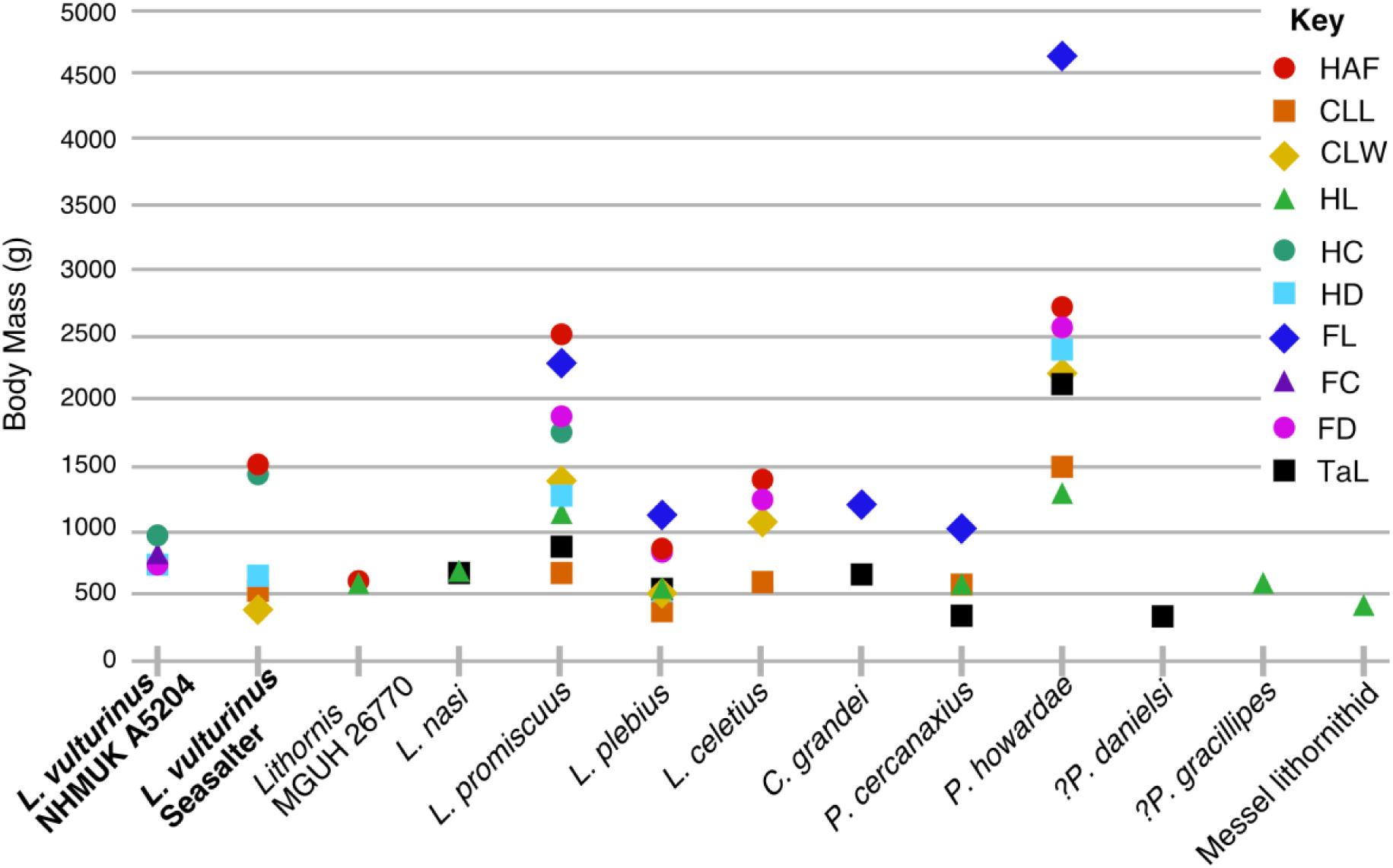
Body mass estimates of selected lithornithids. Body mass correlates are presented as in Benito et al. (2022a), in order of increasing Percent Prediction Error from Field et al. (2013): maximum diameter of the coracoid’s humeral articulation facet (HAF), maximum coracoid lateral length (CLL), least coracoid shaft width (CSW), maximum humerus length (HL), least humerus shaft circumference (HC), least humerus shaft diameter in cranial view (HD), maximum femur length (FL), least femur shaft circumference (FC), least femur shaft diameter in cranial view (FD) and maximum tarsometatarsus length (TaL). Specimens described in this study are highlighted in bold. Femur length and tarsometatarsus length for *L.* cf *grandei*, femur length for *L. promiscuus*, and humeral articular facet, coracoid max lateral length, coracoid least width, and tarsometatarsus length for *P. howardae* represent averages taken from multiple specimens.

All skeletal correlates of body mass indicate that the Seasalter specimen was heavier than both the neotype and MGUH 26770, the two other specimens in our sample that have been referred to *L. vulturinus*. The diameter of the humeral articular facet of the coracoid (HAF) is considered the most reliable single skeletal correlate of body mass in flying crown birds (Field et al., 2013), thus we consider our estimated body mass of the new block to be more accurate than the neotype as the coracoid is preserved. Based on this measurement, the Seasalter specimen yields a mean body mass estimate of 1487 g. The next most accurate metric that can be obtained for both the neotype and the new specimen is the humeral least shaft circumference, which predicts a mean estimate of 956 g for the neotype, and 1423 g for the Seasalter specimen. MGUH 26770 is even smaller, with a HAF measurement yielding a mean body mass estimate of 628 g. This is consistent with the observation that it is slightly smaller in size than the neotype (Bourdon and Lindow, 2015, Mayr and Kitchener, 2025). A fourth specimen referred simply to “*Lithornis*”, NHMUK A5425, has a HAF measurement which yields an intermediate mean body mass estimate of 1144 g. For all individual specimens in our sample for which we could obtain a measurement of the HAF, this measurement yielded the greatest body mass estimate. If this metric is as accurate for lithornithids as it is for birds in the sample of Field et al. (2013), then body mass estimates derived from most other measurements are underestimates.

### Differential diagnosis of *L. vulturinus*

We add the following features to the differential diagnosis of *L. vulturinus* from Bourdon and Lindow (2015) that further distinguish it from other lithornithids. *L. vulturinus* differs from *L. celetius* and *Pseudocrypturus* in having a proximally directed projection on the tip of the lateral process of the coracoid. *L. vulturinus* is distinguished from *Paracathartes*, *Pseudocrypturus*, and *L. plebius* by the hooked shape of the lateral cnemial crest of the tibiotarsus. We tentatively suggest that the absence of a tuber on the proximal part of the minor metacarpal is diagnostic of *L. vulturinus*, though this may be due to weathering of both the neotype and the new block. We have observed this feature in every other known lithornithid taxon, but cannot confirm its presence or absence in MGUH 26770 as the proximal part of the minor metacarpal is obscured by overlying bones. It is present in NHMUK A5425 ‘*Lithornis*’.

### Diagnostic features of the new Seasalter specimen

We identified the new fossil as a lithornithid based on the following combination of features identified by Houde (1988), Stidham et al. (2014), and Bourdon and Lindow (2015) as diagnostic for this clade: the prominent hooked acromion of the scapula, crossed coracoidal sulci of the sternum, a very small dorsal tubercle relative to the size of the ventral tubercle of the humerus, a small concavity between the proximal part of the deltopectoral crest and the distal humeral head in cranial view, the capital incision which forms a notch between ventral tubercle and humeral head in proximal view, the prominent and spherical intercotylar eminence of the tarsometatarsus, and the block-like asulcate hypotarsus lacking hypotarsal canals.

Specific character states that optimise as synapomorphies of Lithornithidae in our analyses that were scorable for this specimen are as follows: the coracoidal sulci are crossed at the midline (C79:0), the posterior margin of the sternum lacks distinct posteriorly projected lateral or medial processes (C83:0), the transverse ligament sulcus of the humerus is deep (C111:0), and the ventral ramus of the ulnare has a tubercle where it joins the dorsal margin (C132:1). We scored ovoid foramina in the lateral surfaces of the centra of the thoracic vertebrae as being present (C67:2); this is extremely difficult to observe in our CT data due to the similarity in radiodensity of the matrix and the fossil but can be seen by the naked eye in thoracic vertebrae visible on the surface of the block.

Unfortunately, the broken deltopectoral crest and poorly defined lateral margins of the sternum in the new specimen impede species level diagnosis somewhat, as these are the only two character traits other than body size listed by Houde (1988) in the diagnosis of *L. vulturinus*. We cannot ascertain the presence of most of the features identified by Bourdon and Lindow (2015) in their amended differential diagnosis of *L. vulturinus* because the relevant bones are missing.

However, their “shallow oblique groove extending from distal part of crista bicipitalis to dorsal margin of sulcus transversus” appears to be present, and the scapular body of the Seasalter fossil is slender and poorly curved, which agrees with their species diagnosis.

Upon adding the new block to the character matrix of Nesbitt and Clarke (2016), we found that it agreed in all 28 scorable characters it shares with the neotype, and in all 25 scorable characters shared with MGUH 26770. Of these characters, three are not plesiomorphic for Lithornithidae: the distalmost end of the scapula is tapered to a fine point (C105:1), the deltopectoral crest is angled steeply relative to the shaft (C116:0), and the proximal portion of the ventral side of metacarpal III is smooth (142:0).

Houde (1988)’s diagnosis of *L. vulturinus* is that it is larger than *L. nasi*, *?L. hookeri*, *L. celetius*, and *L. plebius.* Given that the new specimen’s estimated body mass is greater than that of the neotype, as well as the newly described ?*Pseudocrypturus gracilipes* and ?*Pseudocrypturus danielsi*, it belonging to any of these species is less likely. The new specimen’s estimated body mass is in turn much less than that estimated for *L. promiscuus* and *Paracathartes howardae*. No discernible differences exist between the new specimen and the *L. vulturinus* neotype apart from the slightly larger size of the new fossil, which could reasonably be attributed to intraspecific variation or sexual dimorphism.

### Phylogenetic analysis

To test our referral of the Seasalter fossil to *L. vulturinus*, we incorporated the neotype, the Seasalter fossil, and MGUH 26770 as distinct operational taxonomic units for all analyses. Lithornithidae was recovered as a monophyletic group in all analyses. The neotype and new fossil were consistently recovered in a polytomy with MGUH 26770, *Lithornis promiscuus*, *Lithornis plebius*, *Lithornis celetius*, and *Paracathartes howardae*. Our consistent recovery of the new Seasalter fossil in a polytomy with the neotype and MGUH 26770 lends confidence to our assignment of this specimen to *L. vulturinus*.

### Unconstrained analysis (Figures 22, 23)

Analysis of the Nesbitt and Clarke (2016) dataset under maximum parsimony yielded 4 most parsimonious trees of 438 steps (consistency index = 0.447, retention index = 0.690), and a consensus tree of 452 steps. We recovered a monophyletic Lithornithidae sister to Tinamidae, which collapsed into a polytomy. The monophyly of Lithornithidae is supported by sixteen character states: the lateral side of the pterygoid-palatine articulation area has a distinct fossa with a dorsally bounding ridge (C20:2), a foramen or deep fossa is present on the posterior surface between the ventral condyles and otic process of the quadrate (C38:1), the orbital process of the quadrate is hatchet-shaped and shorter than the length of the quadrate body (C41:1), the articular region of the mandible immediately posterior to the articular facet for the quadrate is expanded posterodorsally as a short process (C60:1), the lateral surfaces of the centra of the thoracic vertebrae have central ovoid foramina (C67:2), the posterior margin of the sternum lacks distinct posteriorly projected lateral or medial processes (C83:0), the n. supracoracoideus appears to pass through the coracoid (C99:0), the transverse ligament sulcus of the humerus is deep (C111:0), and the extensor canal of the tibiotarsus is an emarginated groove lacking an ossified supratendinal bridge (C164:1).

**Figure 22.**
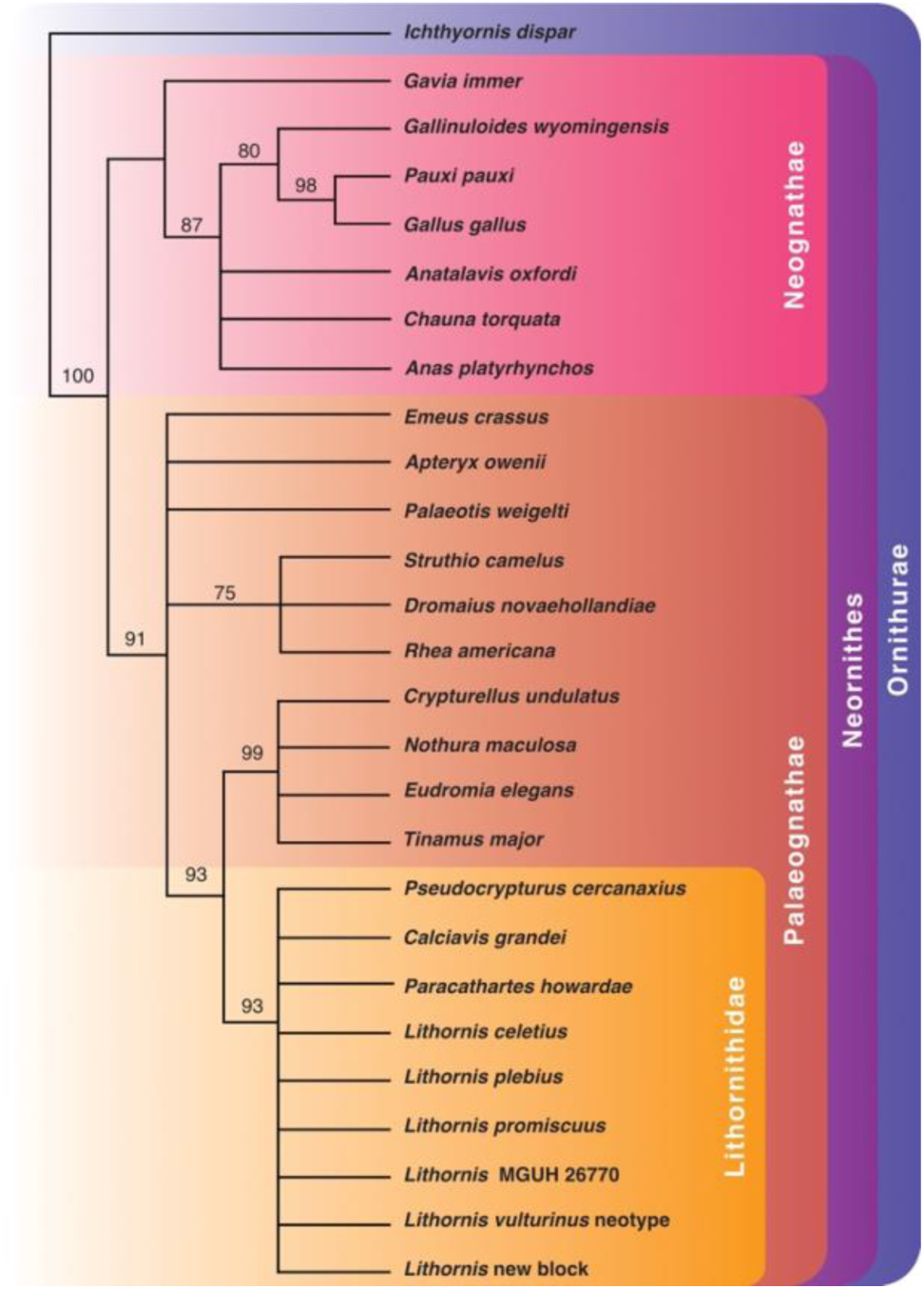
Phylogenetic results of maximum parsimony analysis using a modified version of the morphological matrix from Nesbitt and Clarke (2016) excluding *Apsaravis ukhaana*. Strict consensus of 4 most parsimonious trees with no prior constraints. Bootstrap values are displayed for each node above 50%.

**Figure 23.**
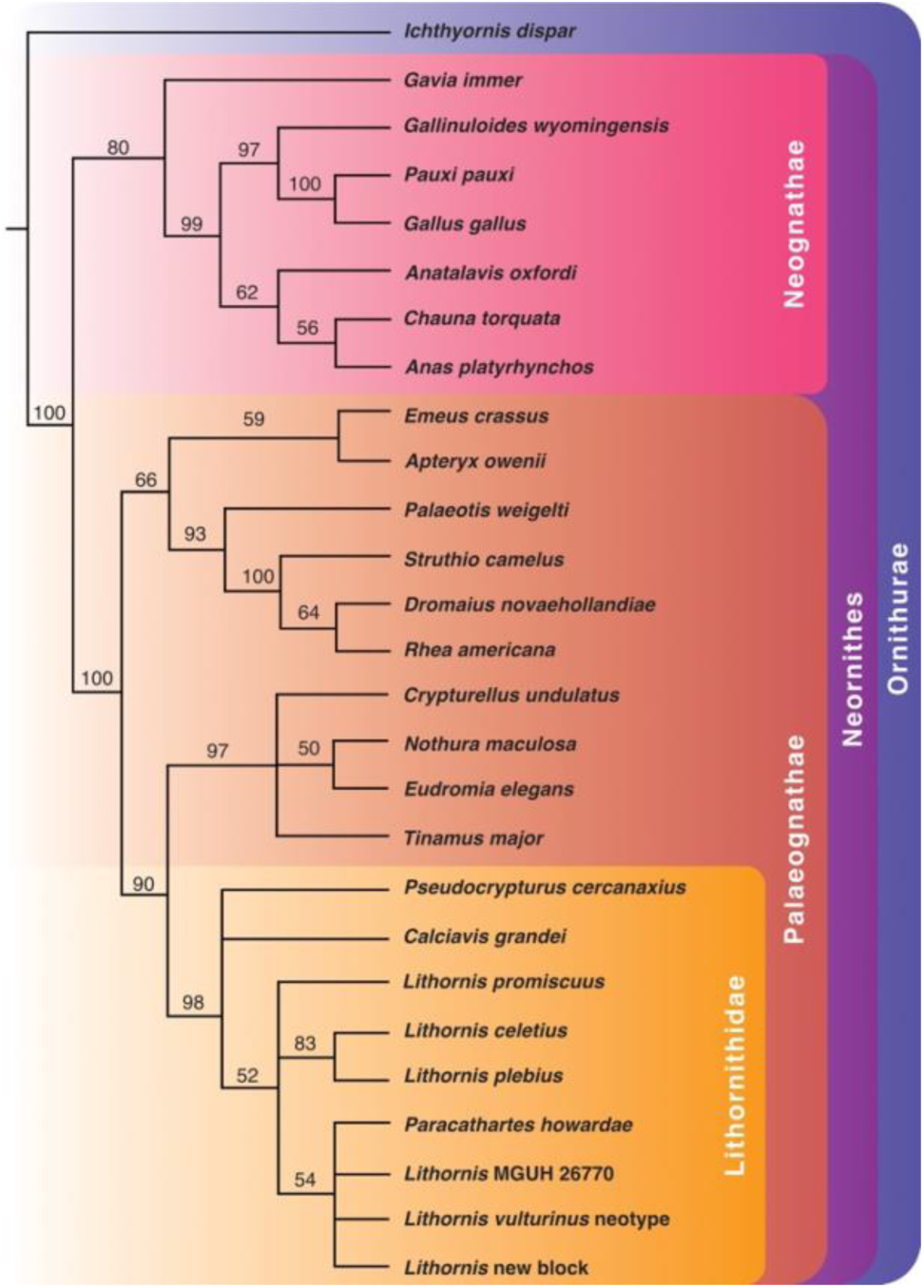
Phylogenetic results from Bayesian inference using a modified version of the morphological matrix from Nesbitt and Clarke (2016) excluding *Apsaravis ukhaana*. No prior constraints were enforced. Node values indicate Bayesian posterior probabilities over 50%.

Constraining the neotype, Seasalter fossil, and MGUH 26770 as a clade required no additional steps. This grouping is supported by a single character: the proximal portion of the ventral side of metacarpal III is smooth (143:0). However, it must be noted that this character is not scorable for MGUH 26770 as the carpometacarpus is preserved in dorsal view, therefore CT scanning of this specimen would be needed to confirm this as an apomorphy of *L. vulturinus*.

Bayesian analysis of the same dataset once again recovered a monophyletic Lithornithidae sister to Tinamidae. The *L. vulturinus* neotype, new block, and MGUH 26770 are recovered in a polytomy with *Paracathartes howardae*, albeit with low bootstrap support, which in turn forms a polytomy with *Lithornis promiscuus* and a *Lithornis celetius* + *Lithornis plebius* clade. *Pseudocrypturus* and *Calciavis* form a polytomy with the clade containing all aforementioned lithornithids. The *L. celetius* + *L. plebius* clade is supported by two synapomorphies: the distal end of the deltopectoral crest of the humerus in dorsal view is at a steep angle relative to the humeral shaft (C116:0), and the proximal portion of the ventral side of the third metacarpal bears a distinct tubercle (C142:1). *Palaeotis weigelti* was found sister to a *Struthio* + *Dromaius* + *Rhea* clade in our Bayesian unconstrained analyses; this result was also found by Nesbitt and Clarke (2016) and by Mayr (2015) in their unconstrained analyses.

### Molecular constraint analysis (Figures 24, 25)

We repeated our maximum parsimony and Bayesian analyses incorporating molecular backbone constraints based on the results of Phillips et al. (2009), Mitchell et al. (2014), Prum et al. (2015), Grealy et al. (2017), Yonezawa et al. (2017), the concatenated dataset of Cloutier et al. (2019), Urantówka et al. (2020), and Almeida et al. (2022). A clade composed of a monophyletic Lithornithidae sister to a monophyletic Tinamidae was recovered in a polytomy with *Dromaius*, *Palaeotis*, *Apteryx*, and *Emeus* in our maximum parsimony analysis, and sister to *Emeus* in our Bayesian analysis. Relationships within Lithornithidae are unresolved in the maximum parsimony analysis, while the Bayesian analysis recovers a *Pseudocrypturus* + *Calciavis* clade and an *L. celetius* + *L. plebius* clade in a polytomy with all other lithornithids.

**Figure 24.**
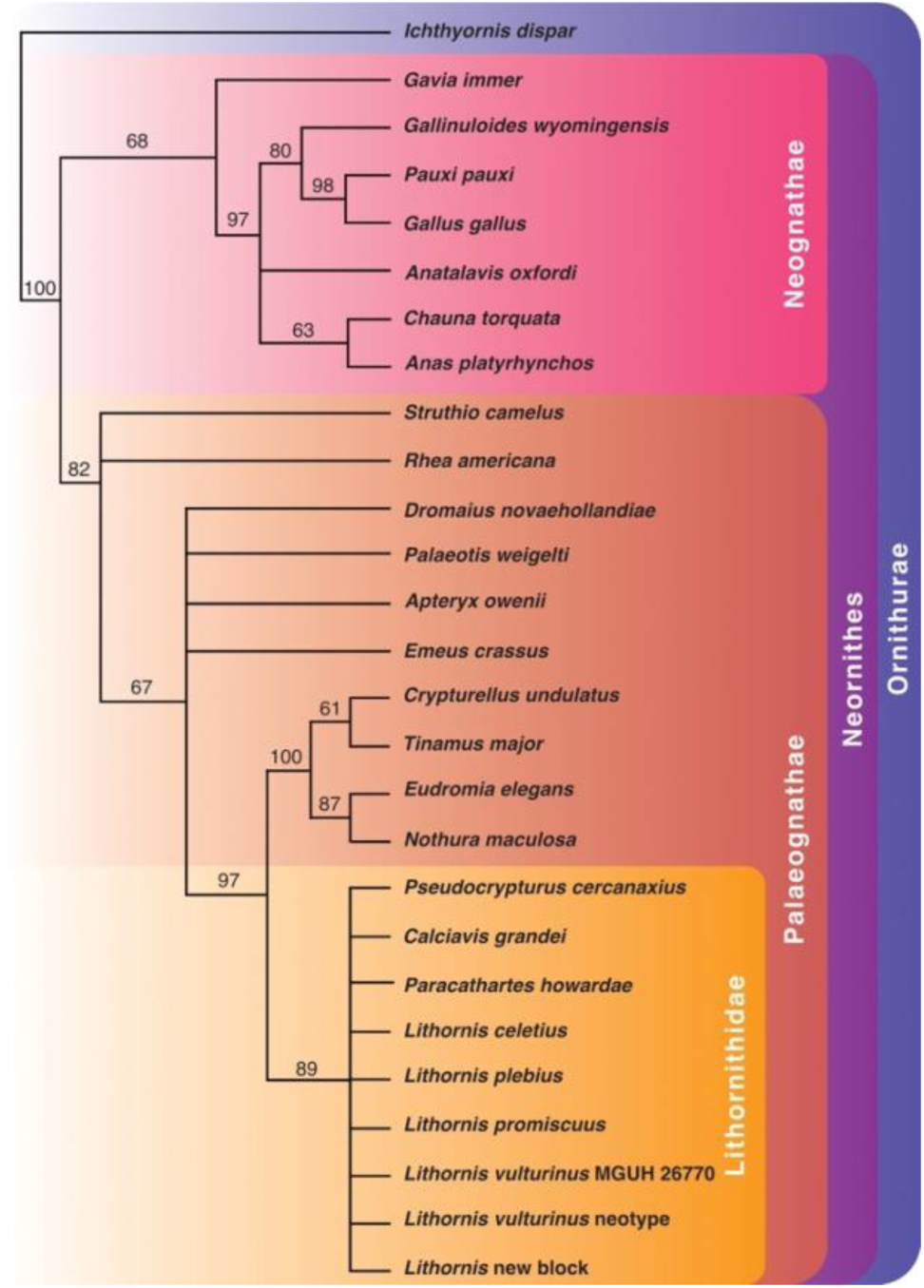
Phylogenetic results of maximum parsimony analysis using a modified version of the morphological matrix from Nesbitt and Clarke (2016) excluding *Apsaravis ukhaana*. Strict consensus of 203 most parsimonious trees with relationships constrained to match the molecular analyses of Phillips et al., 2009; Mitchell et al., 2014; Prum et al., 2015; Grealy et al., 2017; Yonezawa et al., 2017; the concatenated dataset of Cloutier et al., 2019; Urantówka et al., 2020; and Almeida et al., 2021. Bootstrap values are displayed for each node above 50%.

**Figure 25.**
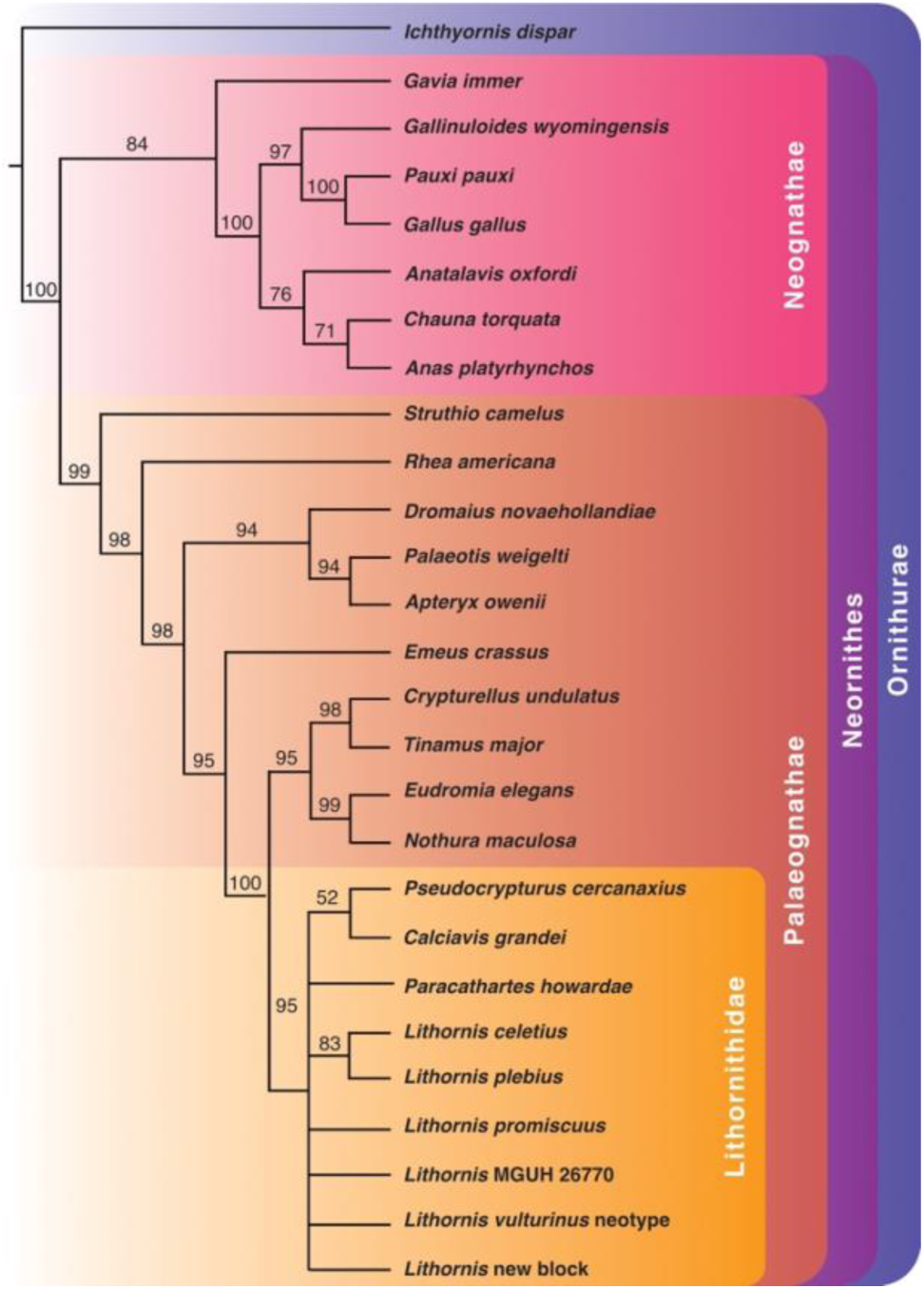
Phylogenetic results from Bayesian inference using a modified version of the morphological matrix from Nesbitt and Clarke (2016) excluding *Apsaravis ukhaana*. Relationships were constrained to match the molecular analyses of Phillips et al., 2009; Mitchell et al., 2014; Prum et al., 2015; Grealy et al., 2017; Yonezawa et al., 2017; the concatenated dataset of Cloutier et al., 2019; Urantówka et al., 2020; and Almeida et al., 2021. Node values indicate Bayesian posterior probabilities over 50%.

### Molecular constraint analysis with lithornithids constrained as the sister taxon to all other palaeognaths (Figures 26, 27)

Under these constraints, all resolution among lithornithids is once again lost under maximum parsimony, with a monophyletic Lithornithidae forming a polytomy that is sister to the remaining palaeognaths. The monophyly of Lithornithidae is supported by twenty character states (C12:0, C20:2, C23:1, C226:0, C48:1, C60:1, C64:1, C67:2, C83:0, C95:1, C103:0, C104:2, C110:1, C111:0, C132:1, C134:1, C140:1,

**Figure 26.**
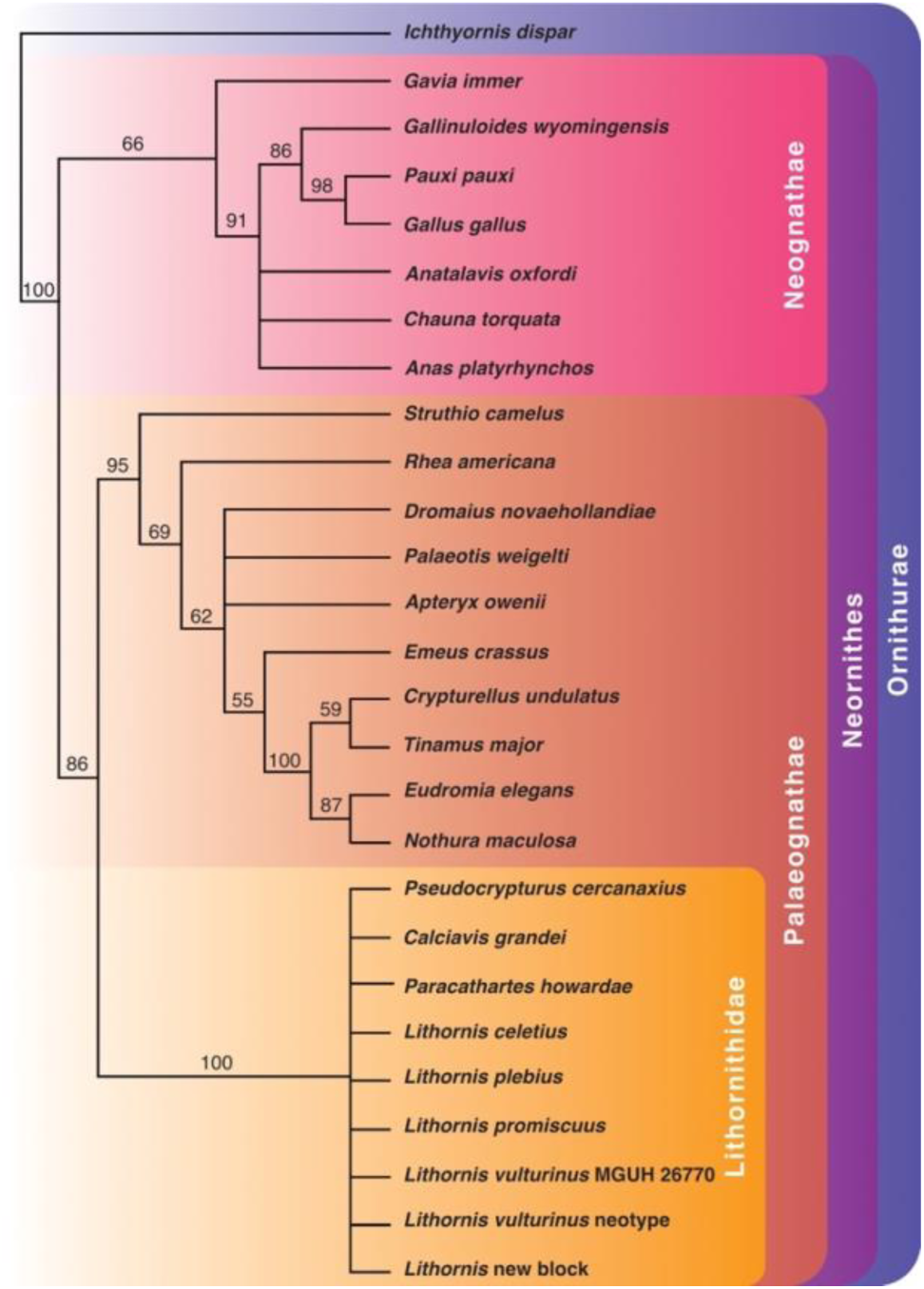
Phylogenetic results of maximum parsimony analysis using a modified version of the morphological matrix from Nesbitt and Clarke (2016) excluding *Apsaravis ukhaana*. Strict consensus of 108 most parsimonious trees with relationships constrained to match the molecular analyses of Phillips et al., 2009; Mitchell et al., 2014; Prum et al., 2015; Grealy et al., 2017; Yonezawa et al., 2017; the concatenated dataset of Cloutier et al., 2019; Urantówka et al., 2020; and Almeida et al., 2021. Lithornithids were also constrained as the sister taxon of all other palaeognaths. Bootstrap values are displayed for each node above 50%.

**Figure 27.**
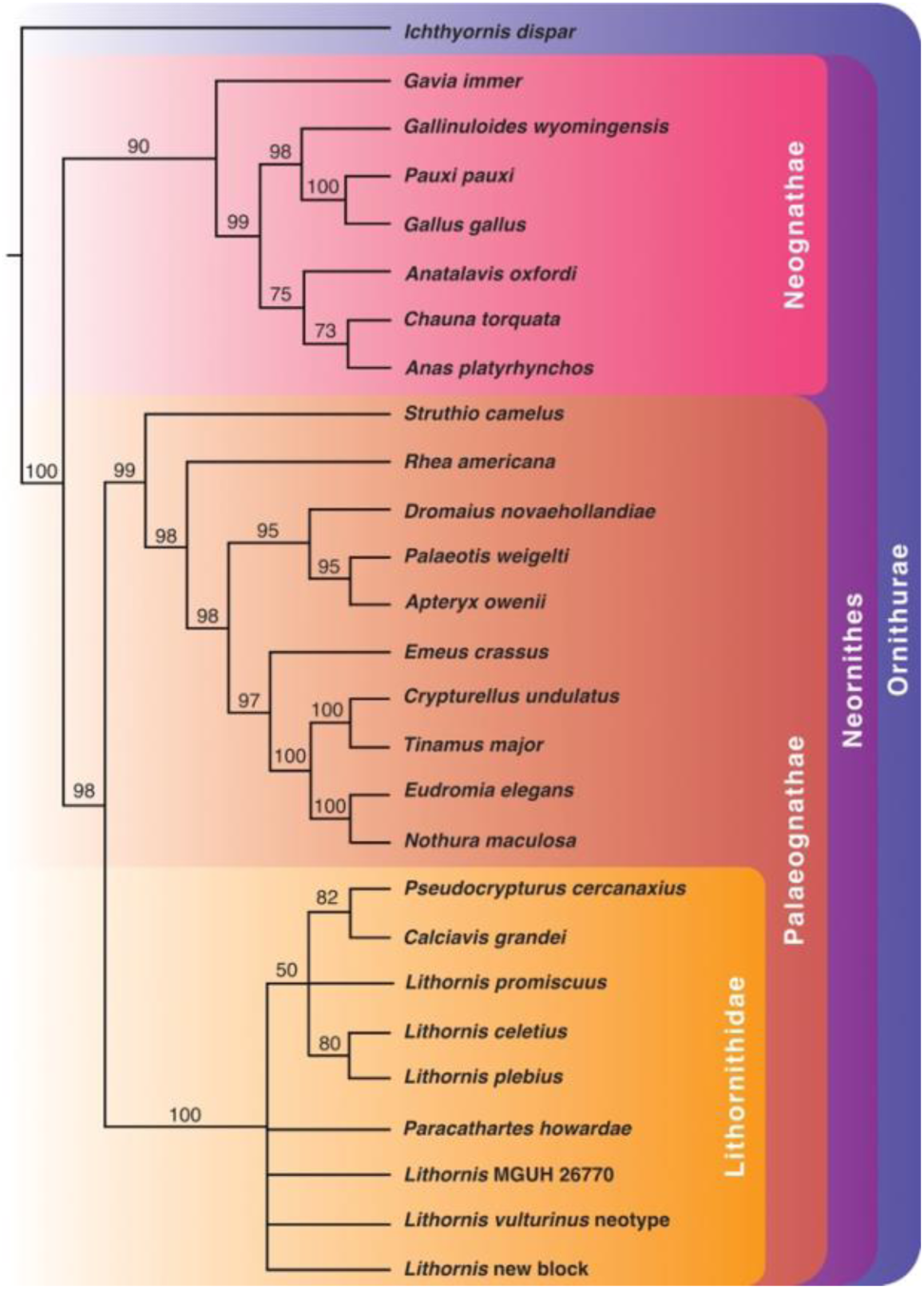
Phylogenetic results from Bayesian inference using a modified version of the morphological matrix from Nesbitt and Clarke (2016) excluding *Apsaravis ukhaana*. Relationships were constrained to match the molecular analyses of Phillips et al., 2009; Mitchell et al., 2014; Prum et al., 2015; Grealy et al., 2017; Yonezawa et al., 2017; the concatenated dataset of Cloutier et al., 2019; Urantówka et al., 2020; and Almeida et al., 2021. Lithornithids were also constrained as the sister taxon of all other palaeognaths. Node values indicate Bayesian posterior probabilities over 50%.

C149:1, C175:1, and C117:0). Of these characters, C79:0 (coracoidal sulci are crossed at the midline) and C104:2 (the acromion of the scapula is laterally hooked with a small foramina on the posterior side) are major visually distinctive features of the lithornithid skeleton. Resolution is also lost within Galloanserae, with *Anatalavis*, *Chauna*, and *Anas* collapsing into a polytomy with Galliformes. Under Bayesian inference, a polytomy consisting of *L. promiscuus*, a *Pseudocrypturus* + *Calciavis* clade, and a *L. celetius* + *L. plebius* clade appears within a polytomy formed by the remaining lithornithids. Resolution is not lost within Galliformes and Anseriformes.

## Discussion

### Referral of Seasalter specimen to *L. vulturinus*

The Seasalter specimen is easily identifiable as a lithornithid, but broad similarities in the postcrania of this group (Houde, 1988) make referral to a specific taxon more difficult. It shares in common all scorable characters from the dataset of Nesbitt and Clarke (2016), but this does not necessarily preclude it belonging to a different species due to the aforementioned similarity in lithornithid postcrania and the lack of a skull in the new specimen. Aside from a somewhat less curved body of the scapula in the Seasalter lithornithid, we could not identify any discernible differences in the shape and form of the bones between this specimen and that of the neotype. The only notable feature that distinguishes the Seasalter specimen from the neotype is its slightly larger size and therefore larger estimated body mass, but this alone is not sufficient to identify it as a separate species as sexual dimorphism is well documented in extant palaeognaths (Olson and Turvey, 2013). Given this information and that the Seasalter fossil was discovered just a few kilometres from the neotype in the same formation, we are confident in taking a conservative approach by assigning the new block to *L. vulturinus* despite its lack of easily diagnosable features.

### Body mass and sexual dimorphism

The large body size of the new specimen compared to the neotype and MGUH 26770 raises the question of possible sexual dimorphism in lithornithids. Reverse sexual dimorphism, in which females are larger than males, is relatively common within palaeognaths (Olson and Turvey, 2013), and is more frequently seen in forest-dwelling palaeognaths rather than those that occupy open habitats (Bunce et al., 2003). Female tinamous tend to be larger than males in nearly all species where sufficient data exists (Winkler et al., 2020). Males of the largest tinamou species, the Gray Tinamou *Tinamus tao*, range between 1325–1863g, with females being larger at 1430–2080g (Cabot et al., 2020a). This reverse sexual dimorphism is especially striking in the Solitary Tinamou *Tinamus solitarius*, where the largest females are nearly twice as heavy as the smallest males (Cabot et al., 2020b). Females had a greater body mass than males in 24 of the 25 tinamou species in the dataset of Tubaro and Bertelli (2003), and were 14% heavier on average. In birds, reverse sexual dimorphism is associated with increased investment in egg laying (Andersson, 1994), and larger size in female tinamous may be the result of selection for laying multiple clutches (Tubaro and Bertelli, 2003).

*L. promiscuus* was named for its presumed polygamous mating behaviour, as fossils were often found associated with large caches of tinamou-like eggshells (Houde, 1988). Houde (1988) interpreted these large caches as the result of simultaneous polygyny and sequential polyandry, in which multiple females lay eggs in the nest of a single male before moving on to find new mates as tinamous do today. Despite this, no evidence of sexual dimorphism in lithornithids has been reported previously (Houde, 1988, Mayr, 2009b, Nesbitt and Clarke, 2016, Mayr and Kitchener, 2025). From our measurements of eight specimens of *Paracathartes howardae* at the USNM, we did not find appreciable differences in estimated body size (Figure 19), though it is not known how many individuals these specimens actually represent.

Houde (1988) considered *Lithornis promiscuus* and *Lithornis plebius* to be separate species on the basis of body size rather than a single sexually dimorphic species, even though the two species are usually found in association. Based on the maximum diameter of the humeral articular facet, we calculated a body mass of 2521 g for *L. promiscuus*, and a body mass of 808 g for *L. plebius*. At over three times the mass of *L. plebius*, this lies outside the range of tinamou sexual dimorphism. Extreme sexual dimorphism existed in the recently extinct moa *Dinornis giganteus* and *Dinornis novaezealandiae* with females nearly three times the mass of males (Bunce et al., 2003), but we consider this scenario unlikely for lithornithids.

Additionally, *L. promiscuus* and *L. plebius* differ in three character states in the Nesbitt and Clarke (2016) matrix. In *L. plebius* the ventral process of the pygostyle is pointed (C75:1) whereas it is club shaped in *L. promiscuus* (C75:2); the deltopectoral crest is angled steeply relative to the shaft in *L. plebius* (C116:0) whereas the angle is shallow in *L. promiscuus* (C116:1); and the anterior portion of the articulation between the ilium and ischium is visible in lateral view in *L. plebius* (C150:0) but not in *L. promiscuus* (C150:1). The differing morphology of the pygostyle was noted by Houde (1988) in his diagnosis of the two species. These taxa did not form an exclusive clade in any of our phylogenetic analyses, nor did they in the analyses of Nesbitt and Clarke (2016).

The new specimen’s estimated mass is nearly a third heavier than the neotype, and more than twice that of MGUH 26770. Should NHMUK A5425 be referable to *L. vulturinus*, this would provide an additional exemplum of a larger specimen belonging to this species. The size difference between the neotype and the new block falls within the range seen between male and female tinamous, whereas that between the new block and MGUH 26770 is strikingly high. We suggest MGUH 26770 be examined histologically to determine if this is indeed an adult individual. Sexual dimorphism remains a possible explanation for this discrepancy, though we are wary of drawing conclusions from so few specimens. Alternatively, Mayr and Kitchener (2025) propose that MGUH 26770 may instead be an exemplar of ?*L. hookeri* or their *L. grandei* (*Calciavis*), which we consider plausible based on our body mass estimates.

### Inferred ecology

Lithornithids have been inferred to be ground-dwelling, probe-feeding birds on the basis of their jaw apparatus apparently permitting distal rhynchokinesis, allowing for prey to be obtained without opening the entire jaw against the substrate (Houde, 1988), as well as evidence for a vibrotactile bill tip organ (du Toit et al., 2020). The asulcate hypotarsus we observe here is consistent with *L. vulturinus* being a ground bird (Mayr, 2016). Among extant birds, this morphology is found in Cathartidae, Sagitariidae, and Cariamiformes (Mayr, 2016), of which the latter two spend considerable time hunting on the ground.

Despite lithornithids possibly spending much of their time foraging on the ground as tinamous do today, there is ample evidence to suggest they were much more proficient in long distance flight than their extant relatives. The sternum is perhaps where the morphologies of these two clades depart most dramatically. The lithornithid sternum lacks the caudal elongation and prominent lateral trabeculae seen in tinamous, both of which provide the large attachment area for the pectoral muscles needed for rapid escape burst flight (Houde, 1988, Widrig et al., 2023).

The humeri of both specimens described here are markedly sigmoid and elongated relative to those of tinamous, which suggests that lithornithids were better adapted for long distance flapping flight with slower wingbeats (Houde, 1988). More definitive conclusions regarding the flight style of lithornithids based on their skeletal morphology will have to await more quantitative ecomorphological comparisons, which are outside the scope of the present study.

### Plesiomorphic neornithine features

Lithornithids bear a variety of features that have been considered plesiomorphic for crown birds (Chen et al., 2025), many of which are evident in these two specimens. For example, the scapular cotyle of the coracoid is circular and deeply excavated, rather than being a flat articular facet. An excavated cotyle is presumably plesiomorphic for Neornithes, and appears in distantly related groups (Mayr, 2021a). The transition from a concave cotyle to a flat articular facet has occurred at least 13 times within Neornithes, including within Palaeognathae, as tinamous have a flat articular facet (C89:1)(Mayr, 2021a).

Another feature that can be observed on both specimens is the clearly crossed coracoidal sulci of the sternum. Sulci crossed at the midline occur in all known lithornithid sterna (Houde, 1988, Nesbitt and Clarke, 2016, Mayr and Kitchener, 2025). This feature is common among early neornithines in the fossil record (Dyke, 2001). It can be observed in the extinct groups Sandcoleiformes (Houde and Olson, 1992) and Presbyornithidae (Dyke, 2001), and the Cenomanian aged *Iaceornis marshi* with a preserved sternum (Clarke, 2004). In *Ichthyornis dispar* specimen YPM 1450 the right sulcus passes below the left for several millimetres (Marsh, 1880, Clarke, 2004), and the sulci are also crossed in YPM 1461 (Clarke, 2004). From our own observations, it can be seen in numerous family level clades widely distributed across extant non-passeriform neoavians. We noted sulci that are slightly crossed at the midline in Steatornithidae, Aramidae, Phoenicopteridae, Burhinidae, Eurypygidae, Phaethontidae, Ciconiidae, Phalacrocoracidae, Threskiornithidae, Scopidae, and Accipitridae. Sulci that are more strongly crossed, reminiscent of the condition in Lithornithidae, occur in Musophagidae, Ardeidae, Pandionidae, Strigidae, Bucconidae, and Falconidae. In their survey of avian pectoral girdle characters, Chen et al. (2025) reported crossed sulci in *Ichthyornis*, *Phoenicopterus*, *Corythaeola*, Grues, *Burhinus*, Phaethontimorphae, Procellariimorphae, *Elanus*, *Leptosumus*, and *Micrastur*. In all birds with crossed sulci, the left sulcus is dorsal to the right, suggesting that this feature had a single evolutionary origin (Clarke, 2004). Based on its distribution among stemward ornithurines such as *Ichthyornis* and early neornithines, it is likely that this feature is also ancestral for crown birds and was lost multiple times within Neornithes in a similar manner to the excavated scapular cotyle. Both the functional significance and developmental mechanisms contributing to crossed coracoidal sulci have not yet been investigated, and it is unknown why multiple reversals to the uncrossed state occurred.

This striking example of nonpathological bilateral skeletal asymmetry is deserving of further research, as such asymmetries are rare among birds (Hatch, 1985). In the Wry-billed Plover *Anarhynchus frontalis* the bill bends to the right (Wiersma and Kirwan, 2023), and in some oystercatchers (genus *Haematopus*) the bill is bent in a minority of individuals, usually to the left (Hockey, 1981). Crossbills in the genus *Loxia* are named for their lower mandibles offset either to the left or right relative to the upper mandible, which the birds use for feeding on conifer seeds (Edelaar et al., 2005). The ‘Akepas and the ‘Akeke’e, Hawaiian honeycreepers of the genus *Loxops*, have crossed mandibles similar to those of crossbills, though the crossing is less conspicuous and involves only the rhamphotheca rather than the underlying skeleton (Richards and Bock, 1973, Hatch, 1985, Lepson and Pratt, 2020). Bilateral asymmetry also occurs in the external ears of some owls, allowing them to more accurately pinpoint sounds (Norberg, 1977). It is notable that these asymmetries all involve jaw or cranial elements, and crossed coracoidal sulci are therefore the only known example of postcranial skeletal asymmetry to occur in birds.

### Phylogenetic analysis

The possibility exists for Lithornithidae to be a paraphyletic assemblage of stem palaeognaths, or a polyphyletic grouping as proposed by Houde (1988), who observed that the bone histology of *Paracathartes* more closely resembled that of extant “ratites” and suggested this may be due to a closer phylogenetic affinity to extant ratites than to other known lithornithids. Though this histological difference is more likely due to the larger body size of *Paracathartes* (Nesbitt and Clarke, 2016), the hypothesis of lithornithid paraphyly or polyphyly cannot be excluded, as stem members of all major extant palaeognath lineages are expected to have been small and volant, similar to the lithornithid morphotype (Almeida et al., 2022, Widrig and Field, 2022). We did not recover any evidence for a paraphyletic or polyphyletic Lithornithidae, though firmly rejecting this scenario will require the discovery and analysis of a greater number of stemward palaeognaths, including identifiable members of extant clades, which are currently lacking in the fossil record (Widrig and Field, 2022).

We did, however, recover limited evidence for subclades within Lithornithidae. The Green River Formation taxa *Calciavis grandei* and *Pseudocrypturus cercanaxius* were recovered as sister taxa in our Bayesian analyses with molecular constraint (posterior probability 52%) and with lithornithids constrained as the sister taxon of all other palaeognaths (posterior probability 82%). They were also found as sister taxa in Nesbitt and Clarke (2016)’s strict consensus with Lithornithidae constrained as the sister taxa of Neornithes and “ratites”, as well as their strict consensus in which *Paracathartes* was constrained as the sister taxon of extant “ratites”. Based on what the authors consider a lack of features that unambiguously differentiate *Calciavis* from every species within the genus *Lithornis*, Mayr and Kitchener (2025) assigned *Calciavis* to *Lithornis*, and refer several of the Walton on the Naze specimens (NMS.Z.2021.40.201, NMS.Z.2021.40.202, and NMS.Z.2021.40.203) to *Lithornis* cf. *grandei*. Despite amending the scorings of two characters for *Calciavis* that Mayr and Kitchener (2025) believed to be erroneously causing *Calciavis* to group with *Pseudocrypturus* (C105 and C117 in our analyses), we nonetheless recovered a Green River clade in two of our Bayesian analyses. Based on our frequent recovery of all lithornithids as a polytomy and never recovering a monophyletic *Lithornis* clade, we consider strong evidence for generic differences in lithornithids to be lacking, and consider the evidence for *Calciavis* being a separate genus to be equivocal.

*Lithornis celetius* and *Lithornis plebius* formed a clade in all Bayesian analyses (bootstrap value 83% with no constraints and with molecular constraints, 80% with molecular constraints and lithornithids constrained as stem palaeognaths). They also appeared as sister taxa in Nesbitt and Clarke (2016)’s strict consensus in which *Paracathartes* was constrained as the sister taxon of extant “ratites”. Of these two relatively small North American lithornithids, *L. celetius* is slightly older; they were not sympatric as in the previous example (Houde, 1988, Widrig and Field, 2022).

We did not recover the sister taxon relationship of *Lithornis promiscuus* and *Paracathartes howardae* that Nesbitt and Clarke (2016) found in their strict consensus when Lithornithidae was constrained as the sister taxon of Neornithes, Palaeognathae, and “ratites”, when *Paracathartes* was constrained as the sister taxon of extant “ratites”, and when relationships were constrained to the molecular trees produced by Phillips et al. (2009), Baker et al. (2014), and Mitchell et al. (2014) in any of our analyses. We additionally found no support for a position of *Pseudocrypturus* outside a clade composed of the remaining lithornithids as found by Yonezawa et al. (2017) in their alternative analysis.

Interestingly, once molecular constraints are added to enforce the sister group relationship between tinamids and dinornithids, lithornithids continue to appear as the sister group to tinamids, with *Emeus crassus* outside the Lithornithidae + Tinamidae clade. This is contrary to the results of Nesbitt and Clarke (2016), who recovered lithornithids as stem palaeognaths when this constraint was added, but is similar to the result obtained by Worthy et al. (2017) in their unweighted maximum parsimony analysis in which a molecular constraint was added. This result seems to be directly related to volancy, as approximately half of the synapomorphies supporting this clade are related to the flight apparatus. However, we acknowledge that such a relationship cannot be refuted on the basis of molecular and current fossil evidence. We support Nesbitt and Clarke (2016)’s conclusion that the position of Lithornithidae as stem palaeognaths remains tentative and requires further investigation.

## Conclusions

We assign the new specimen from Seasalter to *Lithornis vulturinus*, adding a wealth of new material contributing to our understanding of this species. The possibility of sexual size dimorphism within *L. vulturinus* and Lithornithidae as a whole merits further investigation, as this could have implications for ancestral state reconstruction of parental care and egg laying behaviour within Palaeognathae. This will require confirmation of the ontogenetic stage and species referral of MGUH 26770, as it is strikingly small in comparison with the Seasalter specimen. Our phylogenetic analysis achieved limited resolution within Lithornithidae, which is nonetheless recovered as a monophyletic group. The true phylogenetic position of Lithornithidae with respect to the rest of Palaeognathae remains unclear, and likely will not be resolved without the discovery of additional stem palaeognaths. Also meriting further investigation are the evolutionary history and functional implications of crossed coracoidal sulci of the sternum, as this feature of lithornithids likely also appeared in the ancestral crown bird. By providing a more in-depth description of *Lithornis vulturinus* material from the London Clay, we hope to eventually facilitate the identification of other examples of stem palaeognaths from the early Paleocene and Late Cretaceous, in order to eventually clarify the early evolutionary history of Palaeognathae as a whole.

## Collaboration

KEW conceptualised the project, performed digital segmentation, made the figures, performed the phylogenetic analyses, and wrote the first draft. DJF supervised the project. KEW and DJF wrote the final draft. Jack Smith collected the Seasalter fossil and brought it to Cambridge.

## Supplementary Tables

Measurements of postcranial elements of NHMUK A5204 and the Seasalter *Lithornis vulturinus* specimens.

**Table S1.** Measurements of the scapulae of *Lithornis vulturinus* specimens NHMUK A5204 and the Seasalter specimen. Total length is measured from the tip of the acromion process to the furthest distal point on the scapular blade. The glenoid facet length is the maximum craniocaudal length of the glenoid. The omal height corresponds to the maximum dorsoventral extension of the omal end of the scapula, measured as the distance between the dorsalmost point of the omal end to the ventralmost point of the glenoid. Measurements that are unreliable or unrepresentative of true size due to breakage or otherwise incomplete elements are denoted by asterisks (*). Measurements that could not be taken are donated by a dash (-). All measurements are in mm.

| Specimen | Total length | Glenoid facet length | Omal height |
| --- | --- | --- | --- |
| Neotype | 66.87* | 7.40 | 10.99 |
| Seasalter left | 75.96* | 9.33 | 11.54 |
| Seasalter right | 65.51* | 8.97 | 11.49 |

**Table S2.** Measurements of the humeri of *Lithornis vulturinus* specimens NHMUK A5204 and the Seasalter specimen. Total length is the maximum proximodistal length of the humerus. Least circumference, least diameter, and midshaft width are taken at the narrowest portion of the humeral shaft, with midshaft width representing the shortest line between two points on the exterior passing through the centerpoint and least diameter the longest. Proximal craniocaudal width corresponds to the maximum craniocaudal extension of the humeral head, and proximal dorsoventral width is measured as the maximum distance between the dorsal and ventral humeral tubercles, including them. Measurements that are unreliable or unrepresentative of true size due to breakage or otherwise incomplete elements are denoted by asterisks (*). Measurements that could not be taken are denoted by a dash (-). All measurements are in mm.

| Specimen | Total length | Least circumference | Least diameter | Proximal craniocaudal width | Proximal dorsoventral width | Midshaft width |
| --- | --- | --- | --- | --- | --- | --- |
| **NHМУK**<br>**A5204** | 90.63* | 20.94 | 6.31 | 6.54 | 17.82 | 7.08 |
| **Seasalter** | 87.74* | 21.53 | 5.88 | 6.87 | 19.00 | 7.43 |

**Table S3.**
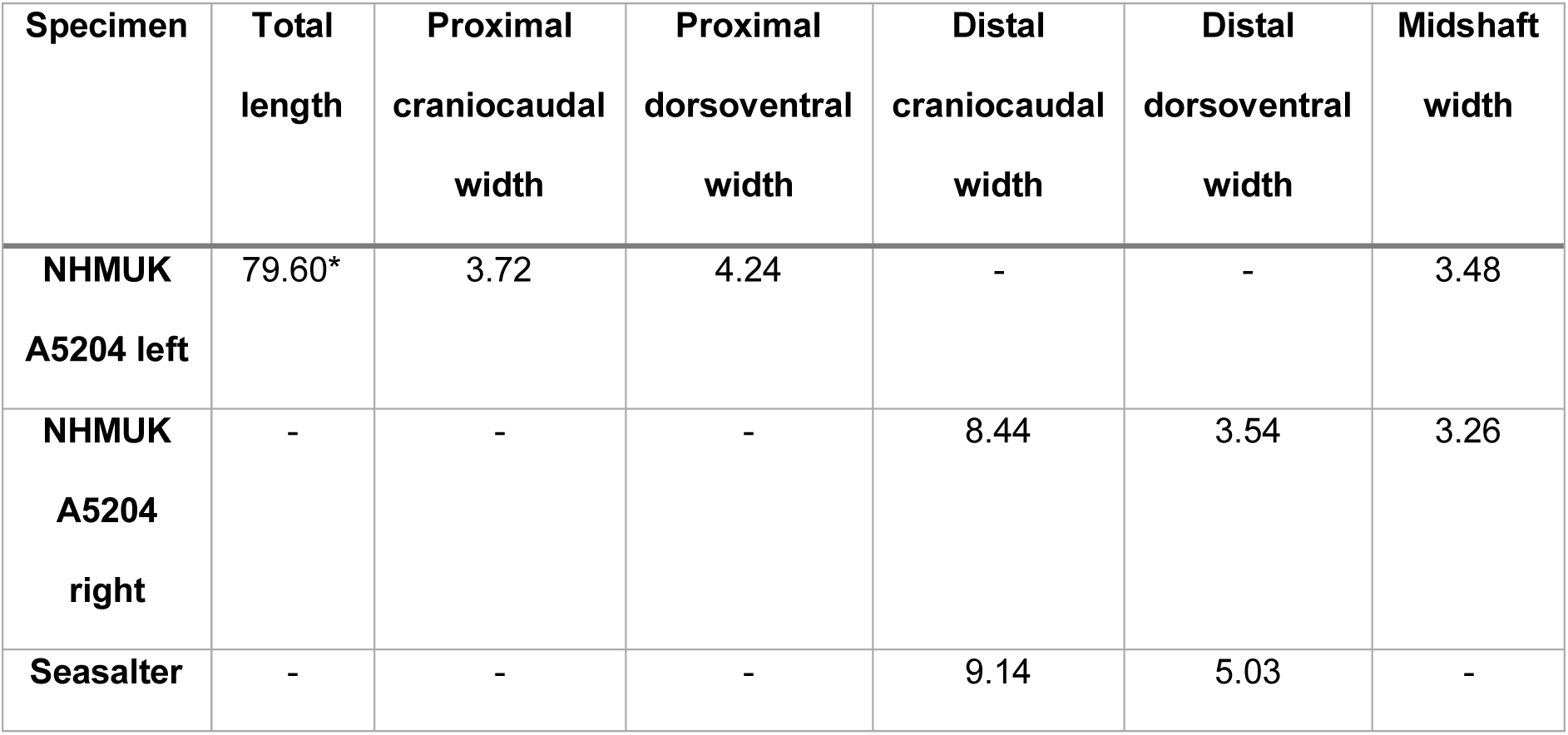
Measurements of the radii of *Lithornis vulturinus* specimens NHMUK A5204 and the Seasalter specimen. Measurements that are unreliable or unrepresentative of true size due to breakage or otherwise incomplete elements are denoted by asterisks (*). Measurements that could not be taken are denoted by a dash (-). All measurements are in mm.

| **Specimen** | **Total length** | **Proximal craniocaudal width** | **Proximal dorsoventral width** | **Distal craniocaudal width** | **Distal dorsoventral width** | **Midshaft width** |
| --- | --- | --- | --- | --- | --- | --- |
| **NHМУK A5204 left** | 79.60* | 3.72 | 4.24 | - | - | 3.48 |
| **NHМУK A5204 right** | - | - | - | 8.44 | 3.54 | 3.26 |
| **Seasalter** | - | - | - | 9.14 | 5.03 | - |

**Table S4.** Measurements of the ulnae of *Lithornis vulturinus* specimens NHMUK A5204 and the Seasalter specimen. Measurements that are unreliable or unrepresentative of true size due to breakage or otherwise incomplete elements are denoted by asterisks (*). Measurements that could not be taken are denoted by a dash (-). All measurements are in mm.

| **Specimen** | **Total length** | **Proximal craniocaudal width** | **Proximal dorsoventral width** | **Distal craniocaudal width** | **Distal dorsoventral width** | **Midshaft width** |
| --- | --- | --- | --- | --- | --- | --- |
| **NHМУK A5204 left** | 83.41* | 10.28 | 8.28 | - | - | 5.32 |
| **NHMUK**<br>**A5204**<br>**right** | 45.46* | - | - | - | - |  |
| **Seasalter** | 32.03* | - | - | 10.63 | 6.80 | 4.47 |

**Table S5.** Measurements of the carpometacarpi of *Lithornis vulturinus* specimens NHMUK A5204 and the Seasalter specimen. Proximal craniocaudal width represents the maximum craniocaudal extension of the proximal carpometacarpus, including the extensor process. The proximal dorsoventral width corresponds to the maximum dorsoventral extension of the proximal carpometacarpus, including the pisiform process. Measurements that are unreliable or unrepresentative of true size due to breakage or otherwise incomplete elements are denoted by asterisks (*). Measurements that could not be taken are denoted by a dash (-). All measurements are in mm.

| **Specimen** | **Total length** | **Proximal**<br>**craniocaudal width** | **Proximal**<br>**dorsoventral width** |
| --- | --- | --- | --- |
| **NHMUK A5204 left** | 22.35* | 9.39* | 5.91 |
| **NHMUK A5204 right** | 15.61* | - | - |
| **Seasalter** | - | - | - |

**Table S6.** Measurements of the sterna of *Lithornis vulturinus* specimens NHMUK A5204 and the Seasalter specimen. Total length represents the maximum craniocaudal distance between the sternal rostrum and the caudal margin. Rostral width, craniolateral process width, and caudal width were all measured on the sternal side that was most complete. Rostral width was measured from the midline of the sternal rostrum to the maximum lateral extension of the coracoid pillar. Craniolateral process width is measured from the sternal midline to the maximum lateral extension of the craniolateral process. Caudal width is the maximum mediolateral width of the caudal margin. Maximum keel depth represents the distance between the cranioproximal edge of the sternal keel to its greatest ventral extension. Measurements that are unreliable or unrepresentative of true size due to breakage or otherwise incomplete elements are denoted by asterisks (*). Measurements that could not be taken are denoted by a dash (-). All measurements are in mm.

| Specimen | Total length | Rostral width | Craniolateral process width | Caudal width | Maximum keel depth |
| --- | --- | --- | --- | --- | --- |
| NHUK<br>A5204 | 67.39 | 10.49 | 22.12 | - | - |
| Seasalter | 77.92 | 10.11 | 21.265* | 30.07* | 22.88 |

**Table S7.** Measurements of the femora of *Lithornis vulturinus* specimen NHMUK A5204. Least circumference, least diameter, and midshaft width are taken at the narrowest portion of the femoral shaft, with midshaft width representing the shortest line between two points on the exterior passing through the centerpoint and least diameter the longest. Proximal mediolateral width includes the femoral head and femoral trochanter. Measurements that are unreliable or unrepresentative of true size due to breakage or otherwise incomplete elements are denoted by asterisks (*). Measurements that could not be taken are denoted by a dash (-). All measurements are in mm.

| Specimen | Total length | Least circumference | Least diameter | Proximal craniocaudal width | Proximal mediolateral width | Distal mediolateral width | Distal craniocaudal width |
| --- | --- | --- | --- | --- | --- | --- | --- |
| NHUK<br>A5204<br>proximal | 18.96<br>* | 18.72 | 5.16 | 7.01 | 12.05 | - | - |
| NHUK<br>A5204<br>distal | 26.83<br>* | 15.39 | 4.61 | - | - | 10.26 | 6.67 |

**Table S8.** Measurements of the tibiotarsi of *Lithornis vulturinus* specimens NHMUK A5204 and the Seasalter specimen. Measurements that are unreliable or unrepresentative of true size due to breakage or otherwise incomplete elements are denoted by asterisks (*). Measurements that could not be taken are denoted by a dash (-). All measurements are in mm.

| Specimen | Total height | Proximal<br>craniocaudal<br>width | Proximal<br>mediolateral<br>width | Midshaft width |
| --- | --- | --- | --- | --- |
| NHMUK A5204<br>left | 13.68* | 10.51 | 13.84 | - |
| NHMUK A5204<br>right | 24.92* | 7.36* | 12.48* | - |
| Seasalter<br>proximal | 39.53* | - | - | - |
| Seasalter shaft | 61.34* | - | - | 5.44 |

**Table S9.** Measurements of the coracoids of the Seasalter *Lithornis vulturinus*. Total length is measured as the maximum length from omal to sternal end. Sternal facet length represents the maximum mediolateral length of the sternal facet of the coracoid. The least coracoid width is measured as the minimum mediolateral width of the coracoid shaft. The glenoid facet diameter represents the maximum diameter of the glenoid (humeral) facet. Measurements that are unreliable or unrepresentative of true size due to breakage or otherwise incomplete elements are denoted by asterisks (*). Measurements that could not be taken are denoted by a dash (-). All measurements are in mm.

| Specimen | Total<br>length | Sternal<br>facet<br>length | Least<br>coracoid<br>width | Glenoid<br>facet<br>diameter | Scapular<br>cotyle max<br>diameter | Lateral<br>process<br>max.<br>height |
| --- | --- | --- | --- | --- | --- | --- |
| **Neotype** | - | - | - | - | - | - |
| **Seasalter left** | 41.85 | 21.45 | 3.19 | 9.46 | 4.58* | 3.78 |
| **Seasalter right** | 41.79 | - | 4.25 | 8.05 | 5.86 | - |

**Table S10.** Measurements of the tarsometatarsi of the Seasalter *Lithornis vulturinus* specimen. Proximal dorsoplantar width includes the hypotarsus. Measurements that are unreliable or unrepresentative of true size due to breakage or otherwise incomplete elements are denoted by asterisks (*). Measurements that could not be taken are denoted by a dash (-). All measurements are in mm.

| **Specimen** | **Total length** | **Proximal dorsoplantar width** | **Proximal mediolateral width** | **Midshaft width** |
| --- | --- | --- | --- | --- |
| **Seasalter left** | 22.86* | 10.08 | 11.97 | - |
| **Seasalter right** | 35.93* | 10.43 | 10.92 | 6.06 |

